# Feeding-fasting cycle of obesogenic food determines glucocorticoid neuromodulation of cortico-hippocampal activities sustaining long-term memory

**DOI:** 10.1101/2025.03.21.644570

**Authors:** Prabahan Chakraborty, Yann Dromard, Emilie M André, Maheva Dedin, Margarita Arango-Lievano, Antoine Besnard, Thamyris Santos Silva, Jean-Christophe Helbling, Guillaume Ferreira, Etienne Challet, Marie-Pierre Moisan, Freddy Jeanneteau

## Abstract

**Background:** Highly caloric food consumed around the clock perturbs the metabolism and cognitive functioning. We hypothesized that obesogenic food could alter neuronal representations of memory depending on the feeding-fasting cycle.

**Methods:** We tracked memory performance, dendritic spine dynamics and neuronal representations of memory in C57Bl6J mice fed obesogenic food *ad libitum* from peri-adolescence. We aimed to correct energy rich diet-induced plasticity deficits and cognitive impairment with time-restricted feeding in males and females. We further used chemogenetics, pharmacology and knock-in mice to investigate functional correlates underlying diet-induced neurocognitive impairments.

**Results:** We find that changes in the feeding-fasting cycle reverted the effects of *ad libitum* obesogenic food on memory impairment in both sexes (n=55, *p*=0.003). Concurrently, it also corrected the increased dendritic spine maintenance and neuroactivity in hippocampus and the decreased spine maintenance and activity in parietal cortex (n=48, *p*<0.005). Bi-directional effects in cortex and hippocampus mediated by glucocorticoid signalling are causal to behavioural changes (n=91, *p*=0.0008), and scaling hippocampal with cortical activities restored memory in mice fed obesogenic food (n=44, *p*=0.02).

**Conclusion:** These results indicate that meal scheduling is a promising approach to confront glucocorticoid signalling bias and memory deficits caused by obesogenic food.

**Research in context:** *Evidence before this study:* What and when we eat contributes to our health. This is particularly worrisome for kids and adolescents because of the lifelong effects that unrestricted snacking on highly caloric food could cause on brain maturation. A variety of school policies and nutritional programs have emerged to prevent poor nutritional habits. But obesity is on the rise and a major cause of neurological disabilities difficult to detect and treat. Human studies are limited by the size and duration of sampling with low resolution metrics to prove causality between nutritional habits and cognitive health trajectory. Animal studies showed that all-day snacking on highly caloric food disrupts innate biological rhythms that influence hormonal secretions, neuronal structure and function in brain regions that encode, store and retrieve memories. It isn’t known if, like adipocytes and hepatocytes, the brain in obesity can develop glucocorticoid resistance -a state that would prevent the robust but complex effects of this hormone on memory- to the point that researchers still question whether glucocorticoids are a cause or solution to obesity related-brain comorbidities.

*Added value of this study:* Longitudinal sampling of several metrics at multiple timepoints in mice fed highly caloric food since peri-adolescence up to adulthood showed that the trajectory of obesity-related brain comorbidities is corrected when reinstating the feeding/fasting cycle, albeit consuming highly caloric food. Glucocorticoid resistance -manifesting as receptor phosphorylation deficits impeding coincidence detection between glucocorticoid and neuronal activities -was reversible when reinstating the feeding/fasting cycle, albeit consuming highly caloric food. Studies in receptor mutant mice lacking a phosphorylation site-independent of glucocorticoids showed it is required to reinstate neuroplasticity to changes of feeding/fasting cycle, albeit consuming highly caloric food. Fos-trapping experiments showed less engagement of pyramidal neurons in the cortex when activity-dependent phosphorylation of glucocorticoid receptor was low, and more in the hippocampus of mice fed obesogenic diet, which reinstating the feeding/fasting cycle reverted, albeit consuming highly caloric food. Finally, chemogenetic experiments confirmed the requirement for the co-engagement of cortical and hippocampal pyramidal neurons to fully remember, despite poor nutritional habits.

*Implications of all the available evidence:* The cortico-hippocampal activities necessary for remembering are uncoupled by obesogenic food consumed *ad libitum* but not on meal scheduling, extending neuroimaging correlation studies in obese adolescents. Poor nutritional habits cause glucocorticoid resistance in the brain as previously suggested, with altered neuronal representation of memory that meal scheduling corrected. This result should transform school policies and familial nutritional habits to promote cognitive health. Future research will develop allosteric ligands targeting phosphorylation motifs in the glucocorticoid receptor as more specific alternative to orthosteric ligands for the treatment of obesity-related brain comorbidities.

## Introduction

Fighting the obesity epidemic is a top priority of health practitioners, given that its prevalence has tripled in the last decade. Direct causes include over-nutrition with energy-dense food, reduced physical activity, and lifestyle pattern changes (1), making it important to identify and correct mechanistic components underlying this global epidemic.

One principal factor that drives such physiological maladaptation is the misalignment of endogenous biological rhythms, which synchronizes activity-sleep and feeding-fasting to the appropriate time of the day-night cycle (2). Repeated disruption of such patterns due to mismatched timing of eating and sleeping increases the risk of obesity and comorbidities like mental illnesses (3). This may arise due to extended periods of indoor artificial lighting causing shorter sleep and longer access to food, particularly Western-style diet (high caloric diet enriched in saturated fat and refined sugars), that can disrupt circadian rhythms at the behavioural, molecular and metabolic levels (4).

Circadian glucocorticoid rhythm, for instance, is disrupted by high-fat diet (HFD) in mice (5), hand-in-hand with a change in feeding rhythms (6). Reducing glucocorticoid levels by treatment with a glucocorticoid synthesis-inhibitor, improves recognition memory in HFD-fed mice (7). Other studies reported dysregulation of hippocampal glucocorticoid-receptor (GR) levels rather than its ligand concentration in a model of obesity-related memory deficits (8). Although glucocorticoids use GR to propagate rapid non-genomic and slow genomic signalling, the impairment of long-term synaptic potentiation from induction when fed HFD suggests that rapid signalling is involved, and possibly via GR phosphorylation as it correlates with its reversal (9). Among the multiple GR phosphorylation sites, serine 134 (S134)- mediated signalling triggers BDNF-dependent positive neuroplasticity of glucocorticoids, learning and memory (10) whereas S226-mediated signalling triggers negative plasticity of glucocorticoids and cognitive impairment (11). The ratio of phosphorylation between S134 and S226 reflects the direction of neuroplasticity and memory performance (12) that could explain bidirectional effects of glucocorticoids in obesity (12). Importantly, the human polymorphism Val66Met in *BDNF* gene disrupts activity-dependent secretion of BDNF, S134-mediated neuroplasticity (15) and glucocorticoid sensitivity (16). Consistently, double transgenic mice carrying the *BDNF* Val66Met genotype and a loss-of-function mutation in S134 impaired activity-dependent neuroplasticity of memory without additive effects, confirming that both pathways interact and that the sole glucocorticoid-dependent sites are insufficient to transduce the full spectrum of GR signalling (15). Therefore, coincidence detection of BDNF and glucocorticoids converging on GR-dependent kinases and phosphatases determines the spectrum of signalling responses (17). Given that GR activation impairs neuroplasticity by suppressing BDNF expression in obese mice (18), we reasoned that HFD could alter the GR phosphorylation ratio that supports structural changes from the glucocorticoid-binding pocket to the docking of signalling effectors (19,20). In agreement, low phosphorylation at S134 while high at S226 in the cortex of patients with Alzheimer’s disease correlated with memory impairment (12). Furthermore, substitution of S134 by an alanine residue while retaining S226 in the APP/PS1 mouse model of Alzheimer’s disease accelerated the progression of cortical amyloidosis and synaptic damage (12). While interventions like environmental enrichment reverted GR-signalling defects in obese mice (8) unlike in *BDNF* Val66Met mice (15), whether GR-phosphorylation ratio underlies memory impairment by HFD remains unknown, and could offer a more specific approach to correcting deficits.

To correct obesity-related diseases, a range of nutritional programs emerged, including caloric restriction (e.g. low-fat food), nutrient choice (e.g. ketogenic diet) and intermittent fasting (e.g. time-restricted feeding, TRF). For instance, most subjects with mild cognitive impairment practicing intermittent fasting reverted to the neurotypical aging category, which was more than in subjects not abiding to the practice (21,22). However, human studies are limited by the acute nature of cognitive testing, the mix of different age-groups and the scarcity of long-term follow-ups investigating the effects of a variety of intermittent fasting styles on cognitive functioning (23). Animal models, by contrast, provide a robust system to investigate long-term effects of intermittent fasting designs on cognition. Beneficial effects of TRF were reported on memory and hippocampal synaptic plasticity in adult rodents fed with normal-diet or HFD (24) known to impair cognition in different age-groups (25,26), diet duration (27), neurodegeneration models (28) and aging-related conditions (29,30). Given that memory retrieval is functionally linked to the maintenance and reactivation of collective synapses between task-allocated neurons (31), we ask whether HFD disrupts such mechanisms by altering glucocorticoids signalling, and if TRF can reverse them.

We addressed these questions using longitudinal *in vivo* imaging of dendritic spines in two brain regions involved in recognition memory–the hippocampus and sensory cortex, that are both involved in early and remote memory retrieval (32). This allowed us to track neuroplastic changes of (i) memory-impairment with peri-adolescent obesogenic diet, (ii) its correction *in the same animal* with TRF in adulthood, and (iii) in mouse carriers of the loss-of-function mutation S134A while retaining S226-mediated signalling. We found that the GR-phosphorylation ratio sustained the bi-directional neuroplasticity between cortex and hippocampus by obesogenic-diet. We additionally used chemogenetics and RU486 infusion to determine the functional role of bi-directional neuroplasticity.

## Methods

### Study approval

Experiments abide to the Directive by the Council of European Communities (86/609/EEC) and were approved by the French Ministry of research and ethics committee (CCEA-APAFIS1769-28536, 1574-18707). All efforts were made to minimize animal suffering and reduce their number.

### Animals

Age/weight/sex-matched B6.Cg-*Tg(Thy1-YFP)HJrs*/J, *Fostm2.1(icre/ERT2)^Luo^*/J, B6.Cg-Gt(ROSA)26Sor^tm14(CAG-tdTomato)Hze^/J from Jackson labs, and NR3C1 knock-in mutant Ser134Ala/Ser267Ala (*B6.Tg(Nr3c1^tm2/Jean^)/J)* (15) were group-housed under standard pathogen-free laboratory conditions (12/12 light/dark cycle, lights on at 7 am, 22°C, 60% humidity, food and water *ad libitum*) according to ARRIVE guidelines. See Supplemental information, Table S1 for reagent details.

### *Ad libitum* high fat and sucrose diet (HFD) and time-restricted feeding (TRF)

Mice were fed since weaning (day 21) to either a high fat-high sucrose diet (HF: 4.7 kcal/g, cholesterol 0.195 mg/g (45% kcal saturated fat), 35% carbohydrates with 50% sucrose, D12451 Research Diet) or a control normal chow (NC: 3.3 kcal/g, 8% kcal fat, 45% proteins, 73% carbohydrates, A04 SAFE) for 12 weeks, provided either *ad libitum* or with time-restriction (food access from zt11 to zt1, active period) for the last 4 weeks, an adaption from published protocols (33,34). Here, the effects of 4-weeks TRF examined in young adults superimpose with the *ad libitum* consumption of HF or NC during peri-adolescence. This model did not alter daily calorie intake, energy expenditure, respiratory exchange ratio, circulating glucocorticoid levels or locomotion but increased fat pads over lean mass and caused cognitive impairment (35).

### Thinned skull cortical window, and intra-hippocampal optical window

For the cortex, skull bone in *Thy1*-YFP mice was thinned to transparency using disposable ophthalmic surgical blades (Surgistar, Vista, CA, USA) with skull-implanted head plate over the window for imaging. For the intra-hippocampal hippocampal window, a 3–4 mm craniotomy (AP -2.3 mm, ML 2 mm) was prepared by drilling with permanent flow of cold HEPES-buffered artificial cerebrospinal fluid (aCSF in mM, 120 NaCl, 3.5 KCl, 0.4 KH2PO4, 15 glucose, 1.2 CaCl2, 5 NaHCO3, 1.2 Na2SO4, 20 HEPES, pH=7.4) and removal of dura and cortex by vacuum suction down to callosal projections (36). A detailed map of the pial vasculature and dendritic territories were taken for subsequent relocation as previously described (37).

### Stereotaxic surgery

AAV2-m*Camk2a*-hM3D(Gq)_mCherry-WPRE-hGHp(A), AAV2-m*Camk2a*-hM4D(Gi)_mCherry-WPRE-hGHp(A) (500 nl, Univ Zurich, Switzerland) were used in this study. Eleven weeks-old C57BL6J mice were injected at 2 sites in S1 (AP -0.1 mm and -0.9 mm, ML +/- 2.5 mm, DV -1.0 mm) and/or at one site in dCA1 (AP -1.8 mm, ML +/-1.5 mm, DV -1.3 mm). Multi-sites expression of transgenes was verified post-mortem.

### Genetic tagging of cells

Transgenic mice with *Fos* promoter that drives the expression of c-Fos and ER^T2^-iCre knocked-in the gene’s 3’-UTR (38) were used. Hydroxytamoxifen (4OH-TAM: I.P. 25mg/kg dissolved in 15% Dimethyl-formamide, 85% sesame oil) sends ER^T2^-iCre into the nucleus expressing tdTomato via loxP recombination.

### Statistics

Animal were randomly allocated to experimental groups and analysed by scientists blinded to the groups. There was no a priori calculation of sample size because the number of mice with correct genotype was never exact. No data was excluded. Missing timepoints is due to unexpected death from repeated anaesthesia or lack of testing. Normality of datasets was tested with Shapiro-Wilk test and homogeneity of variance with Spearman’s or Levene’s test. ANOVA was used for multifactorial comparisons with post-hoc analyses reporting significant difference of the means (2-way model: main effects of diet and schedule or DREADD and CNO, 3-way model: main effects of time, diet and schedule or DREADD Gi, Gq and CNO, or genotype, diet and regimen or RU486, vehicle and diet, and their interactions with Prism 9 (GraphPad). We used R for 4-way ANOVA model that included the sex factor and 5-way model that included sex with all other factors in the mutant experiment. The use of the chi-square distribution instead of the F-distribution is justified by the lack of homoscedasticity. For data that did not meet the assumption of parametric analysis, we used the Friedman’s test to determine difference between all factors and repeated measures with post-hoc Wilcoxon’s test and Kruskal-Wallis test for pairwise comparisons. *P* values were corrected for false discovery rate (α=0.05), and Cohen’s *d* calculated as an indicator of effect size between groups (see **supplemental Table S2** for details).

### Role of funders

Funders had no role in study design, data collection, analyses, interpretation, and writing.

## Results

### Time-restricted feeding on HFD ameliorated memory impairments

To explore the impact of TRF on HFD (45% kcal from fat, 17% sucrose) *ad libitum* since weaning, access to food was limited to zt11-to-zt1 (active period) for postnatal week 12-15, while NC-fed controls too stayed either in TRF schedule or *ad libitum*. Several outcome measures were collected at weaning (baseline, week 1), while on HFD before TRF (week 8), and after 4 weeks of TRF (week 12) (**Fig. 1a**). After 8 weeks of HFD, males and females had significant change in body weight (*p*=0.0008, **Supplementary Fig. S1a**), normal locomotor activity (**Supplementary Fig. S1b**), no anxiety (**Supplementary Fig. S1c**), and were thus randomly assigned to the *ad libitum* or TRF groups. At weeks 1, 8 and 12, memory performance was assessed with NOR test (**Fig. 1b**). Both males and females went from normal performance at weaning (week 1) to cognitive impairment when fed HFD *ad libitum* for 8 and 12 weeks (**Fig. 1c**). ANOVA with post-hoc comparisons showed significant difference between group-means (*p*<0.0001) with large effect size (*d*>2) for HFD compared to NC, with no sex-difference **(Supplementary Table S2)**. Remarkably, TRF on HFD reversed memory impairments in both males (*p*=0.02) and females (*p*=0.04) with large effect sizes (male: *d*=1.78, female: *d*=1.9), and TRF on NC had no effect on memory-performance (males: *p*=0.9, *d*=0.7, females: *p=*0.9, *d*=0.6).

**Fig. 1.**
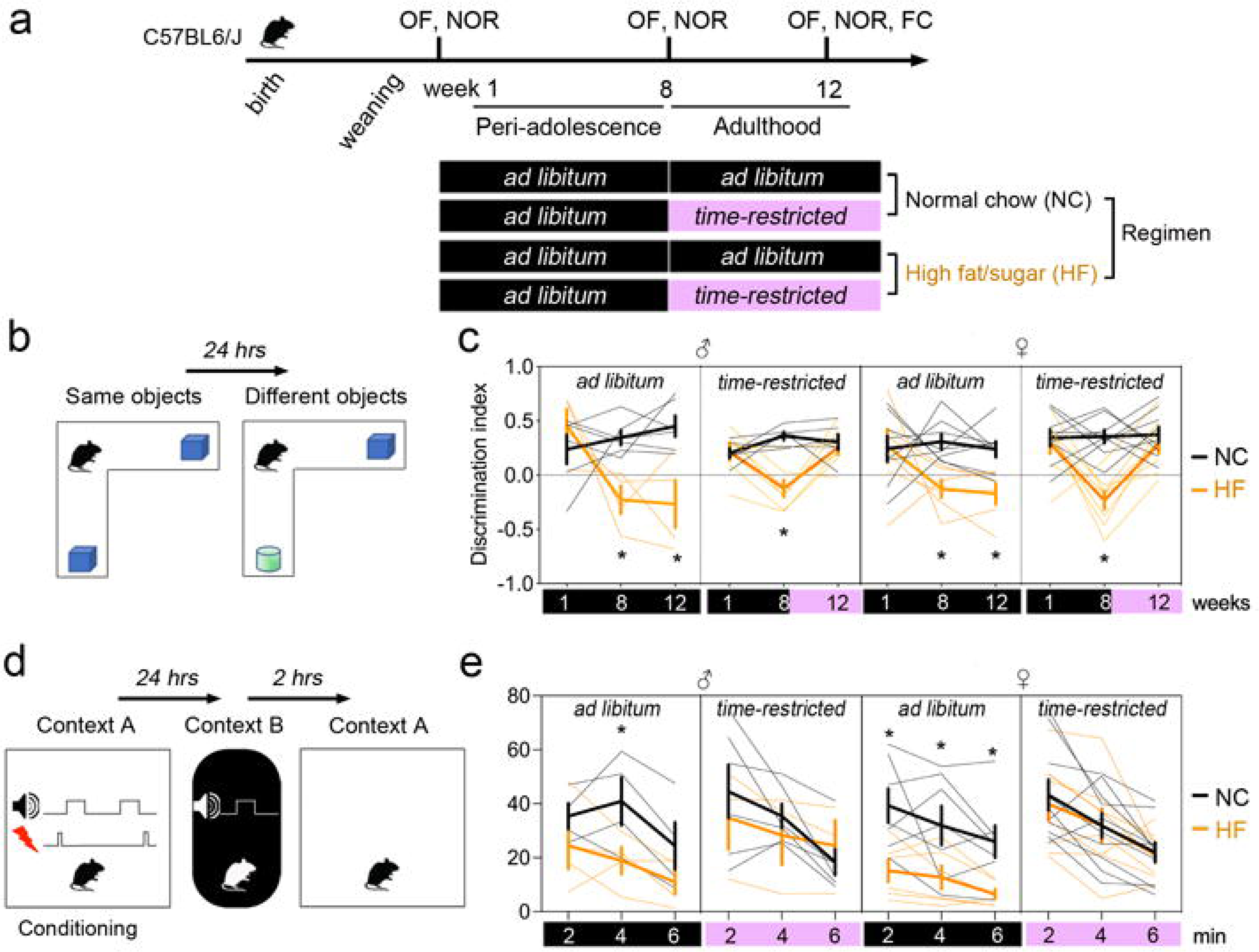
Time-restricted feeding on HFD reverted memory impairments. **a.** Males (n= 23) and females (n = 32) fed high fat/sugar (HF) or normal chow (NC) were subjected to repeat testing in the open field (OF), for novel object recognition memory (NOR) and fear conditioning memory (FC). Black colour: ad libitum, pink colour: time restricted feeding. **b.** Recall of object recognition memory was assessed 24 hrs after learning session with the same objects. **c.** Discrimination of the new object over the old one (index = new/(new+old)). Mice fed NC or HF *ad libitum* (black) or on *restricted-time* (purple). Bold lines indicate means ± SEM, thin lines individual subjects (n = 14 mice with NC *ad libitum*, 11 with HF *ad libitum*, 15 with NC *restricted*, 15 with HF *restricted)*. Four-way ANOVA: Effect of sex x diet x schedule x objects *F*_(2,92)_ = 0.8 *p* = 0.4. Consolidated 3-way ANOVA without sex factor: Effect of diet x schedule x objects *F*_(2,151)_ = 6.08 *p* = 0.002, effect of schedule x objects *F*_(2,151)_ = 5 *p* = 0.007, effect of diet x objects *F*(2,151) = 17.8 *p* < 0.0001, effect of diet x schedule *F*(1,151) = 1.62 *p* = 0.02, effect of diet *F*(1,151) = 48.4 *p* < 0.0001, effect of objects *F*(2,151) = 9.1 *p* = 0.0002, effect of schedule *F*(1,151) = 4.49 *p* = 0.03. See **table S2** for post-hoc analysis \**p* < 0.05 and Cohen’s *d* effect size. **d.** Males (n = 17) and females (n = 31) fed HF or NC were tested for FC memory in adulthood. Electrical foot shocks were delivered during conditioning in context A, while an unpaired tone was delivered in a novel context B. Recall of FC memory was assessed 24 hrs later in context B with the unpaired tone and 2 hrs later in the conditioning context A without shocks. **e.** % Freezing behaviour during recall of the context A. Mice fed NC or HF *ad libitum* (black) on *restricted-time* (purple). Bold lines indicate means ± SEM, thin lines individual subjects (n=10 mice with NC *ad libitum*, 10 with HF *ad libitum*, 13 with NC *restricted*, 12 with HF *restricted).* Chisq analysis: Effect of sex *χ2*(1,40) = 0.15 *p* = 0.69, diet *χ2*(1,40) = 6.5 *p* = 0.01, schedule *χ2*(1,40) = 3.7 *p* = 0.05, time *χ2*(2,40) = 72 *p* < 0.0001, diet x schedule *χ2*(1,40) = 5 *p* = 0.02, time x schedule *χ2*(2,80) = 4.8 *p* = 0.09, diet x time *χ2*(2,80) = 1.4 *p* = 0.4, diet x schedule x time *χ2*(2,80) = 0.4 *p* = 0.7, sex x diet *χ2*(1,40) = 0.03 *p* = 0.8, sex x schedule *χ2*(1,40) = 0.2 *p* = 0.6, sex x time *χ2*(2,80) = 1.1 *p* = 0.5, sex x diet x schedule *χ2*(1,40) = 0.2 *p* = 0.6, sex x schedule x time *χ2*(2,80) = 0.4 *p* = 0.7, sex x diet x time *χ2*(2,80) = 2 *p* = 0.3, sex x diet x schedule x time *χ2*(2,80) = 0.4 *p* = 0.4. See **table S2** for post-hoc analysis \**p* < 0.05 and effect size.

We further validated this finding by testing contextual fear memory in the same mice on week 13 (**Fig. 1d**), as previous studies suggested diet-effects on emotional memory (39). In agreement, *ad libitum* HFD reduced the time of freezing in the conditioning context A as compared to NC-fed mice, without sex-differences (**Fig. 1e, Supplementary Table S2**). ANOVA revealed significant difference between groups (*p*=0.0002) with large effect size (*d*=1.35) specific to shock-paired context A, but not to shock-unpaired tone presentation in context B (*p*=0.9, *d*=0.4). TRF on HFD also reverted emotional memory impairments across sex (males: *p*=0.05, *d*=1.5, females *p*=0.002, *d=*1.7), with no effect on NC-fed animals (males: *p*=0.5, *d*=0.5, females: *p*=0.5, *d*=0.2). Therefore, *ad libitum* HFD degraded both aversive and non-aversive memories, which could be successfully reverted by TRF across sex. Given these results, further experiments were conducted in a mix of both sexes.

### Time-restricted feeding on HFD reversed cortical and hippocampal spine changes

To investigate the dynamics of neuronal alterations accompanying memory restoration on TRF, we imaged *in vivo* remodelling of dendritic spines at the same three timepoints as described above (**Fig. 2a).** Comparisons between the 1^st^ and 2^nd^ images (week 8 vs. 12) indicated dynamic remodelling of dendritic spines due to TRF, while the persistence of this change was recorded in session 3 (week 12 vs. 13) (**Fig. 2b**). Previous studies on the same HFD model indicated that obesogenic diet during adolescence can *enhance* and *reduce* neuronal functions depending on the brain regions (39–41). This prompted us to prepare optical imaging windows atop the sensory cortex (S1) and dorsal hippocampus (dCA1) as both activate during retrieval of episodic memory (32) in transgenic *Thy1*-YFP mice labelling sparse layer-5 pyramidal neurons with bright YFP-fluorescence. Mice use S1 layer-5 pyramidal neurons for detailed exploration (*e.g.* object texture-sensing) (42) and dCA1 pyramidal neurons to form representation of tactile information while encoding episodic memory (43).

**Fig. 2.**
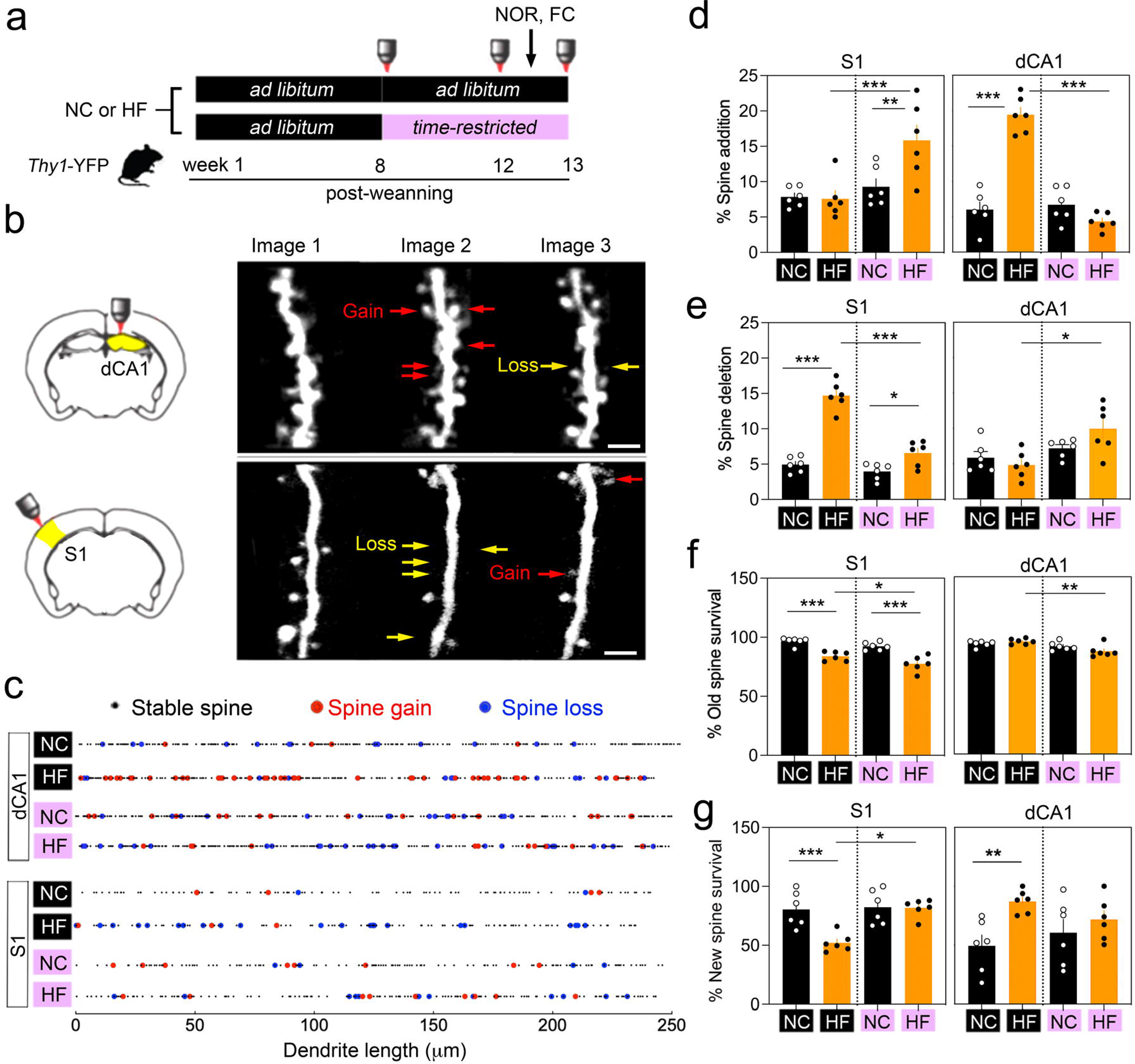
Time-restricted feeding on HFD reverted dendritic spine impairments. **a.** Design with imaging at 3 time points in *Thy1*-YFP males (n = 24) fed with high fat/sugar (HF) or normal chow (NC). Black colour: ad libitum, pink colour: time restricted feeding. **b.** Longitudinal in vivo multi-photon imaging of pyramidal neurons in somatosensory cortex S1 via a transcranial window and in dorsal hippocampus subfield dCA1 via an intrahippocampal window implanted on week 4. Arrows point to dynamic spines. Scale = 4 μm. **c.** Representation of stable (small black dots) and dynamic (red dots for gains, blue dots for losses) spines along dendritic territories between week 8 and 12. Dynamic remodelling comes in clusters defined as 2 or more dendritic spines within 5 μm distance from each other. **d.** % Spine addition between week 8 and 12 (means ± SEM of n = 6 mice/group). Two-way ANOVA For S1: Effect of schedule *F*(1,20) = 12.3 *p* = 0.002, diet *F*(1,20) = 5.2 *p* = 0.03, interaction *F*(1,20) = 6.1 *p* = 0.02. For CA1: Effect of schedule *F*(1,20) = 59 *p* < 0.0001, diet *F*(1,20) = 35 *p* < 0.0001, interaction *F*(1,20) = 71 *p* < 0.0001. See **table S2** for post-hoc analysis \*\**p* < 0.01, \*\*\**p* < 0.001 and effect size. **e.** % Spine elimination between week 8 and 12 (means ± SEM of n = 6 mice/group). Two-way ANOVA For S1: Effect of schedule *F*(1,20) = 52 *p* < 0.0001, diet *F*(1,20) = 95 *p* < 0.0001, interaction *F*(1,20) = 32 *p* < 0.0001. For CA1: Effect of schedule *F*(1,20) = 11 *p* = 0.003, diet *F*(1,20) = 0.8 *p* = 0.3, interaction *F*(1,20) = 3.9 *p* = 0.06. See **table S2** for post-hoc analysis \**p* < 0.05, \*\*\**p* < 0.001 and effect size. **f.** % Old spine maintenance between week 8 and 12 (means ± SEM of n = 6 mice/group). Two-way ANOVA For S1: Effect of schedule *F*(1,20) = 8 *p* < 0.0001, diet *F*(1,20) = 57 *p* < 0.0001, interaction *F*(1,20) = 0.3 *p* = 0.5. For CA1: Effect of schedule *F*(1,20) = 12 *p* = 0.001, diet *F*(1,20) = 0.5 *p* = 0.4, interaction *F*(1,20) = 4.5 *p* = 0.04. See **table S2** for post-hoc analysis \**p* < 0.05, \*\**p* < 0.01, \*\*\**p* < 0.001 and effect size. **g.** % New spine maintenance between week 8 and 13 (means ± SEM of n = 6 mice/group). Two-way ANOVA For S1: Effect of schedule *F*(1,20) = 10 *p* = 0.003, diet *F*(1,20) = 9 *p* = 0.006, interaction *F*(1,20) = 8 *p* = 0.008. For CA1: Effect of schedule *F*(1,20) = 0.05 *p* = 0.8, diet *F*(1,20) = 8 *p* = 0.008, interaction *F*(1,20) = 2.4 *p* = 0.13. See **table S2** for post-hoc analysis \**p* < 0.05, \*\**p* < 0.01, \*\*\**p* < 0.001 and effect size.

A comparison between imaging sessions 1 and 2 showed robust remodelling of dendritic spines often occurring in clusters within 5 μm distance in both S1 and dCA1 (**Fig. 2c**). The extent of changes was exceptionally high and region-specific in mice fed HFD. In particular, only dCA1 neurons increased spine *addition* (**Fig. 2d**) whereas only S1 neurons increased spine *elimination* in mice fed HFD *ad libitum* (**Fig. 2e**). ANOVA showed significant group-mean differences for spine additions (*p*<0.0001) with a large effect size (*d*=5.1) in dCA1 but not in S1 (*p*=0.8, *d*=0.1). Spine eliminations, on the other hand, showed no difference between groups in dCA1 (*p*=0.4, *d*=0.4), with only a significant effect in S1 (*p*<0.0001, *d*=6). The maintenance of pre-existing spines decreased in S1 significantly (*p*<0.0001, *d*=3.5) without being affected in dCA1 (*p*=0.3, *d*=0.8) (**Fig. 2f**), while newly-formed spine survival decreased in S1 (*p*=0.004, *d*=2.3) but increased in dCA1 (*p*=0.004, *d*=2.1) (**Fig. 2g**).

Strikingly, TRF on HFD reversed all these effects with significant difference between groups with large effect sizes **(Supplementary table S2)**. Specifically, TRF improved spine addition, lowered the extent of spine deletion and rescued new spine survival in S1 (**Fig. 2d,e,g**), while it prevented new spine additions and survival in dCA1 (**Fig. 2d,g**). The only exception was that TRF failed to change the reduced maintenance of pre-existing spines in S1 (*p*<0.0001, *d*=2.8) (**Fig. 2f**), despite the increase of spine additions (*p*=0.003, *d*=1.5) (**Fig. 2d**). Remarkably, the normalized dynamics of spine remodelling persisted in the 3^rd^ imaging sessions of S1 and dCA1 on TRF while it remained altered on HFD *ad libitum* (**Supplementary Fig. S2**). Altogether, the results indicate that TRF corrected structural alterations of pyramidal neurons caused by HFD in S1 and dCA1.

### Time-restricted feeding on HFD reverted cortical and hippocampal activity changes

To confirm differential engagement of neurons in S1 vs. dCA1 suggested by dendritic spine remodelling, we harvested brains 1 hour after NOR recall to quantify cFos immediate early gene induction in *Thy1*-YFP pyramidal neurons (**Fig. 3a**). The number of Fos-positive YFP-neurons decreased in S1 and increased in dCA1 with *ad libitum* HFD (**Fig. 3b**), with YFP-neuron proportion within Fos-activated cells only changing in dCA1 (**Fig. 3c**). There were significant differences between groups in dCA1 and S1 with large effect sizes (dCA1: *p*=0.005, *d*=1.7, S1: *p*=0.005, *d*=1.55). Hence, pyramidal neurons were contrastingly engaged between S1 (*less*) and dCA1 (*more*) after recall, and remarkably, TRF reversed these effects (**Fig. 3b**). HFD-fed mice on TRF tend to show less activation in dCA1 (*p*=0.07, *d*=1.7) and significantly more in S1 (*p*=0.0005, *d*=2.12), also decreasing the proportion of YFP neurons within the population of Fos-activated cells in dCA1 (*p*=0.0001, *d*=1.88) (**Fig. 3c**).

**Fig. 3.**
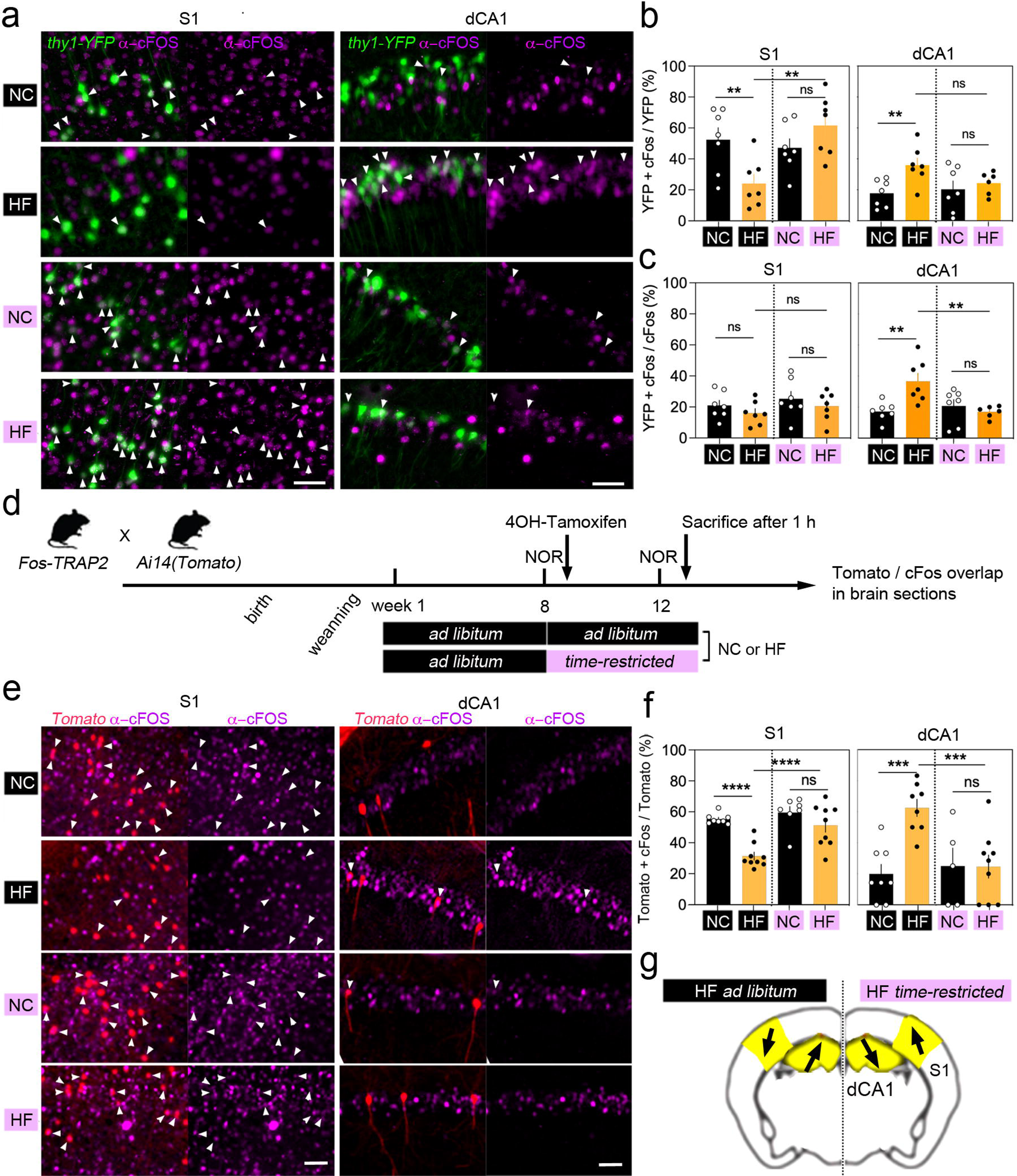
Time-restricted feeding on HFD reverted neuronal engagement deficits. **a.** c-Fos induction labelled with antibodies in pyramidal neurons of somatosensory cortex S1 and dorsal hippocampus subfield dCA1 in *Thy1*-YFP males (n = 28). Arrowheads point to dual labelled cells. HF: high fat/sugar, NC: normal chow. Black colour: ad libitum, pink colour: time restricted feeding. Scale = 50 µm. **b.** % Fos-activated YFP neurons 1 hr after NOR on week 13 (means ± SEM of n = 7 mice/group). Two-way ANOVA for S1: Effect of schedule *F*(1,24) = 4.7 *p* = 0.04, diet *F*(1,24) = 0.6 *p* = 0.4, interaction *F*(1,24) = 12 *p* = 0.001. Two-way ANOVA for CA1 (n = 7 mice/group except n = 6 HF *restricted)*: Effect of schedule *F*(1,23) = 1 *p* = 0.3, diet *F*(1,23) = 6.7 *p* = 0.01, interaction *F*(1,23) = 2.7 *p* = 0.1. See **table S2** for post-hoc analysis \*\**p* < 0.01 and effect size. **c.** % YFP neurons among Fos-activated cells 1 hr after NOR on week 13 (means ± SEM of n = 7 mice/group). Two-way ANOVA for S1: Effect of schedule *F*(1,24) = 1.4 *p* = 0.2, diet *F*(1,24) = 1.7 *p* = 0.1, interaction *F*(1,24) = 0.006 *p* = 0.9. Two-way ANOVA for CA1 (n = 7 mice/group except n = 6 HF *restricted)*: Effect of schedule *F*(1,23) = 4.4 *p* = 0.04, diet *F*(1,23) = 4.6 *p* = 0.04, interaction *F*(1,23) = 9.7 *p* = 0.005 post-hoc analysis \*\**p* < 0.01. See **table S2** for post-hoc statistics and effect size. **d.** Timeline for the genetic tagging in *Fos*-TRAP2;Ai14 males (n = 33) of Fos-activated neurons (tdTomato) engaged by novel object recognition (NOR) testing on week 8 and reactivated (cFos induction) on week 13. **e.** NOR-evoked Fos-induction on week 13 in neurons of S1 and dCA1 engaged by NOR (tagged with tdTomato) on week 8. Arrowheads point to dual labelled cells. Scale = 50 µm. **f.** % Fos-activated tdTomato cells 1 hr after NOR on week 13 (means ± SEM). Two-way ANOVA for S1 (n = 8 mice with NC *ad libitum*, 9 with HF *ad libitum*, 7 with NC *restricted*, 9 with HF *restricted)*: Effect of schedule *F*(1,29) = 12 *p* = 0.001, diet *F*(1,29) = 22 *p* < 0.0001, interaction *F*(1,29) = 5.4 *p* = 0.02. Two-way ANOVA for CA1 (n = 8 mice with NC *ad libitum*, 8 with HF *ad libitum*, 5 with NC *restricted*, 9 with HF *restricted*): Effect of schedule *F*(1,26) = 4.6 *p* = 0.04, diet *F*(1,26) = 7.7 *p* = 0.009, interaction *F*(1,26) = 8 *p* = 0.008. See **table S2** for post-hoc analysis \*\*\**p* < 0.001, analysis \*\*\*\**p* < 0.0001 and effect size. **g.** Summary: HFD enhanced dCA1 activation, decreased S1 activation, and TRF reverted it.

Does HFD also affect re-activation of memory-encoding cells? To address this, cFos-positive cells were marked with a permanent genetic tracer in double-transgenic mice (Floxed-tdTomato crossed with *Fos*-TRAP2(ER^T2^-CRE). These animals were fed HFD *ad libitum* from week 1-to-8, and TRF initiated at adulthood from week 8-to-12. Tomato-tagging was induced by hydroxytamoxifen (4OH-TAM) injection immediately after NOR recall and before schedule change on week 8. Four weeks later during TRF, mice were retested in NOR (with the same objects and test conditions) and brains harvested 1 hour later (**Fig. 3d**). Mice fed HFD *ad libitum* showed lesser re-engagement of NOR memory-encoding cells in S1 (Tomato^+^/cFos^+^ cells *p*<0.0001, *d*=3.8) and greater reactivation in dCA1 (*p*=0.0003, *d*=2.5) (**Fig. 3e**). Remarkably, TRF corrected the engagement of Tomato^+^ cells in dCA1 (*p*=0.0007, *d*=1.9) and in S1 (*p*=0.0001, *d*=1.7) (**Fig. 3e**). Together, the results indicate that TRF on HFD re-engaged memory-activated cells *more* in S1 and *less* in dCA1.

### Corticohippocampal chemogenetic manipulation corrected HFD-induced memory deficits

We next used chemogenetics to determine causality between neuronal engagement and memory performance. In particular, we tested if reversing the contrasting patterns of change in S1 versus dCA1 can correct memory in HFD-fed mice. Animals were bilaterally injected with Designer-Receptor-Exclusively-Activated-by-Designer-Drugs (DREADD) to activate (S1) or inhibit (dCA1) pyramidal neurons, targeted under a *Camk2a* promoter. The DREADD ligand CNO (or vehicle) was injected 45 minutes prior to NOR recall on week 12 (**Fig. 4a**).

**Fig. 4.**
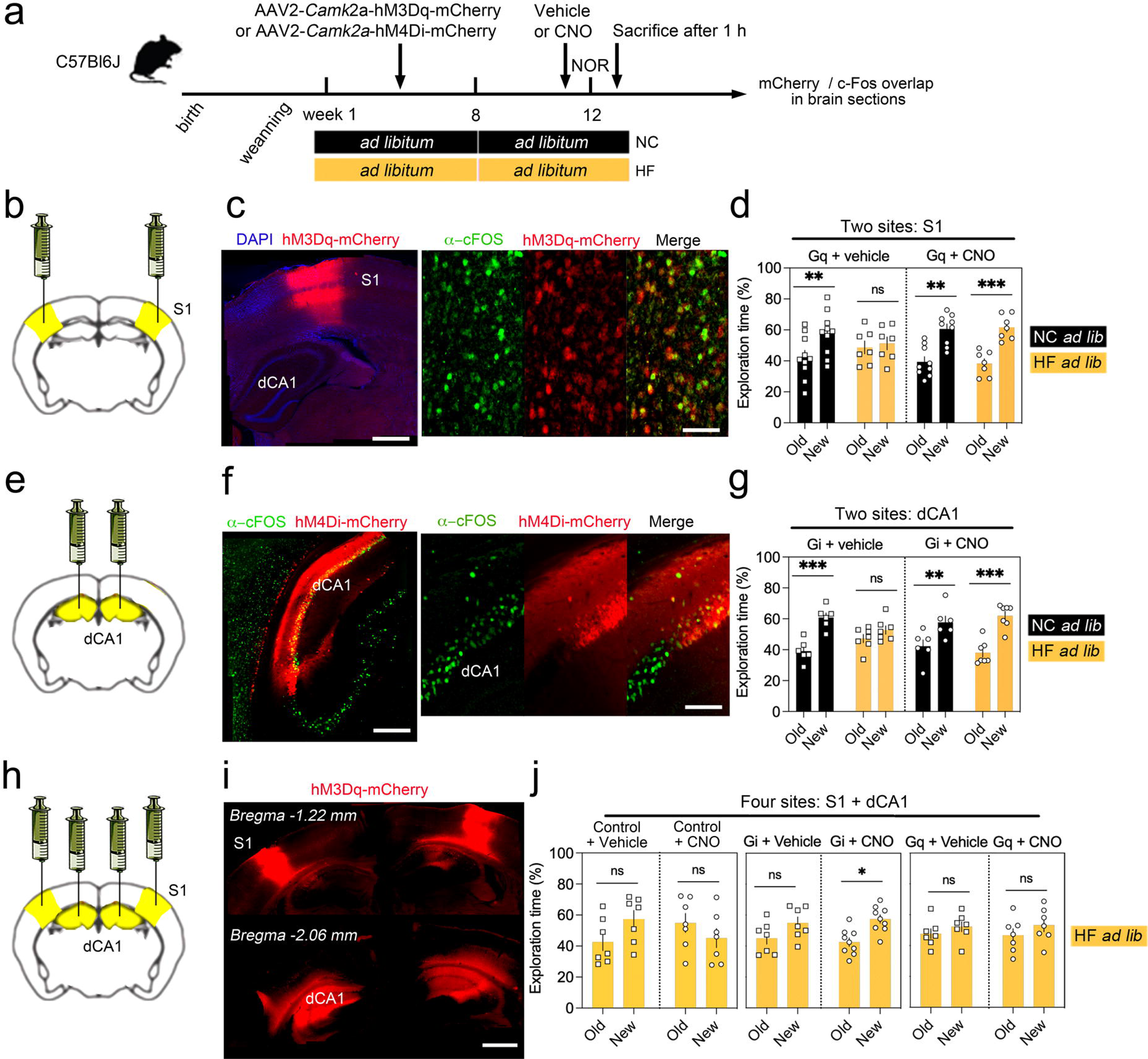
Downplaying dCA1 activity trumps S1 deficits and memory impairment in mice fed obesogenic food. **a.** Timeline. Clozapine-N-oxide (CNO as DREADD ligand) injected I.P. at 2 mg/kg 45 minutes before novel object recognition (NOR) testing. HC: high fat/sugar, NC: normal chow. **b.** Bilateral injections of hM3dq (DREADD activator: Gq) virus in somatosensory cortex S1 in males (n = 24) and females (n = 8). **c.** Transgene expression and CNO-evoked induction of c-Fos labelled with antibodies. Scale = 500 and 40 µm, respectively. **d.** % Time exploring the old versus new objects in the NOR on week 12 (means ± SEM of n = 10 mice with NC *ad libitum+vehicle*, 7 with HF *ad libitum+vehicle*, 9 with NC *ad libitum+CNO*, 7 with HF *ad libitum+CNO)*. Three-way ANOVA: Effect of diet x CNO x objects *F*(1,29) = 0.8 *p* = 0.3, effect of CNO x objects *F*(1,29) = 2.8 *p* = 0.1, effect of diet x objects *F*(1,29) = 0.4 *p* = 0.5, effect of objects *F*(1,29) = 15 *p* = 0.0005, effect of CNO *F*(1,29) = 54 *p* < 0.0001. See **table S2** for post-hoc analysis \*\**p* < 0.01, \*\*\**p* < 0.001 and effect size. **e.** Bilateral injections of hM4di (DREADD inhibitor: Gi) virus in dorsal hippocampus subfield dCA1 in males (n = 16) and females (n = 10). **f.** Transgene expression and CNO-evoked repression of NOR-induced c-Fos labelled with antibodies. Scale = 250 and 65 µm, respectively. **g.** % Time exploring the old versus new objects in the NOR on week 12 (means ± SEM of n = 6 mice with NC *ad libitum+vehicle*, 6 with HF *ad libitum+vehicle*, 7 with NC *ad libitum+CNO*, 7 with HF *ad libitum+CNO)*. Three-way ANOVA: Effect of diet x CNO x objects *F*(1,22) = 3.8 *p* = 0.06, effect of CNO x objects *F*(1,22) = 331 *p* < 0.0001, effect of diet x objects *F*(1,22) = 0.3 *p* = 0.5, effect of objects *F*(1,22) = 28 *p* < 0.0001, effect of CNO *F*(1,22) = 80 *p* < 0.0001. See **table S2** for post-hoc analysis \*\**p* < 0.01, \*\*\**p* < 0.001 and effect size. **h.** Bilateral injections of hM4di (or hM3dq) virus in dCA1 and S1 in males (n = 20) and females (n = 22). **i.** Transgene expression at 4 sites. Scale = 700 µm. **j.** % Time exploring the old versus new objects in the NOR on week 12 (means ± SEM of n = 7 Sham controls with HF *ad libitum+vehicle*, 7 sham controls with HF *ad libitum+CNO*, 7 Gi mice with HF *ad libitum+vehicle*, 9 Gi with HF *ad libitum+CNO*, 7 Gq with HF *ad libitum+vehicle*, 7 Gq with HF *ad libitum+CNO)*. Two-way ANOVA: Effect of objects *F*(1,12) = 0.09 *p* = 0.7, CNO *F*(1,12) = 3.6 *p* = 0.078, interaction *F*(1,12) = 2 *p* = 0.1. Three-way ANOVA: Effect of DREADD x CNO x objects *F*(1,26) = 0.04 *p* = 0.8, effect of CNO x objects *F*(1,26) = 0.24 *p* = 0.6, effect of DREADD x objects *F*(1,26) = 0.9 *p* = 0.3, effect of objects *F*(1,26) = 6.5 *p* = 0.016, effect of DREADD *F*(1,26) = 224 *p* < 0.0001. See **table S2** for post-hoc analysis \**p* < 0.05 and effect size. Gi: DREADD inhibitor, Gq: DREADD activator.

In S1, hM3Dq-activation with CNO **(Fig. 4b)** induced cFos in infected cells (**Fig. 4c**, **Supplementary Fig. 3**), indicating increased engagement of S1 pyramidal neurons. HFD-fed mice improved object discrimination comparable to NC-fed controls when injected with CNO (*p*=0.009, *d*=2.9) unlike with vehicle (*p*=0.7, *d*=0.23) (**Fig. 4d**).

In dCA1, on the other hand, we expressed the DREADD inhibitor hM4Di in pyramidal neurons (**Fig. 4e**), which decreased Fos-induction after CNO injection (**Fig. 4f**, **Supplementary Fig. 4**). HFD-fed mice improved object discrimination comparable to NC-fed controls when injected with CNO (*p*=0.0007, *d*=3.1) unlike with vehicle (*p*=0.3, *d*=0.7) (**Fig. 4g**). Therefore, bi-directional effects of HFD on pyramidal neurons of S1 (deactivation) and dCA1 (hyperactivation) are causal to memory impairment, and reversible with chemogenetics.

### Scaling hippocampal and cortical activities corrected HFD-induced memory deficits

Next, we targeted DREADD expression in both S1 and dCA1 of the same mice to dissect their functional uncoupling in HFD-induced memory deficits (**Fig. 4h**). First, hM4Di-mediated inhibition in both S1 and dCA1 significantly improved memory (**Fig. 4j**, CNO: *p*=0.02, *d*=1.7, Vehicle: *p*=0.18, *d*=0.9) compared to sham-injected, HFD-fed controls. This contrasted with hM3Dq-mediated activation of both S1 and dCA1 that failed to ameliorate memory performance in *ad libitum* HFD-fed mice (**Fig. 4j**, CNO: *p*=0.3, *d*=0.5, Vehicle: *p*=0.5, *d*=0.5). This indicated that dCA1 activation occluded the beneficial effects of S1 activation in the HFD-model, and that down-scaling activity in dCA1pyramidal neurons was necessary and sufficient to correct memory retrieval.

### Blocking GR recapitulates opposite effects in S1 and dCA1 by HFD *ad libitum*

To investigate the underlying mechanism, we injected the GR antagonist RU486, well-known modulator of dendritic spine remodelling and object recognition memory (44,45). We tracked the remodelling of dendritic spines in S1 and dCA1 when RU486 was injected immediately after NOR training (**Fig. 5a**) as it impaired memory retention 24 hours later (*p*=0.01, **Fig. 5b**). Imaging 7-day spine turnover on week 12 (**Fig. 5C**) showed that RU486 inverted the ratio of additions versus eliminations compared to vehicle-injected controls, by enhancing net loss of spines in S1 and gains in dCA1 (*p*=0.03, **Fig. 5d**), similar to the effect of HFD *ad libitum* (**Fig. 2c,d**). We harvested brains 1 hour after NOR for histological examination of GR signalling markers dependent on glucocorticoids (pS226), or BDNF (pS134) (46) (**Fig. 5e**). Strikingly, pS134 levels decreased in S1 but increased in dCA1 of HFD-fed mice, while TRF reversed these opposite effects (**Fig. 5f**) without changing GR levels (**Supplementary Fig. 5**). In contrast, TRF failed to revert HFD-induced changes of pS226 (**Fig. 5g**). These results suggest that specific components of GR signalling could be downregulated in S1 and upregulated in dCA1 by HFD *ad libitum*.

**Fig. 5.**
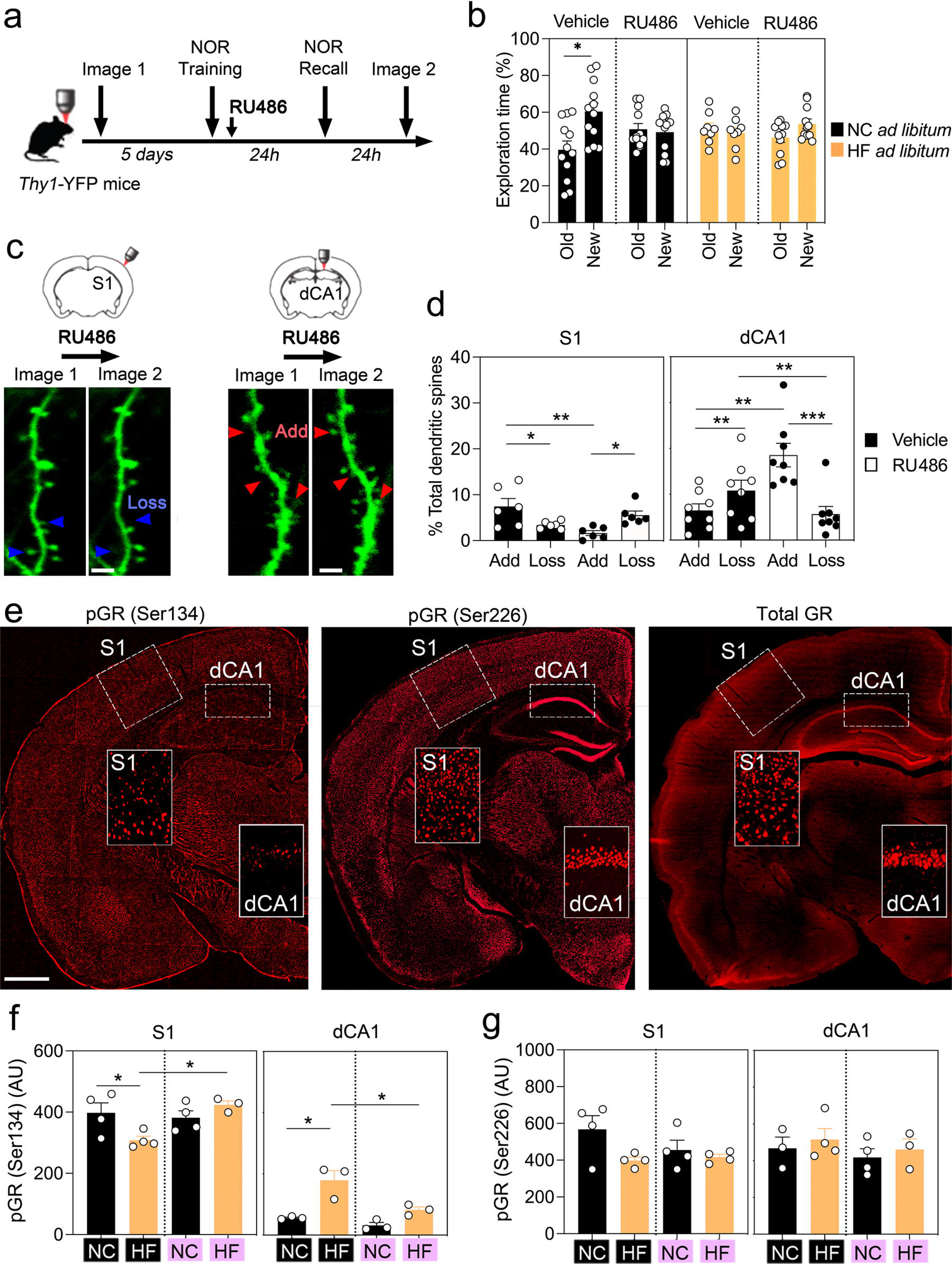
HFD-induced memory deficits associated with GR phosphorylation and function. **a.** Timeline in males (n = 43) during week 12: RU486 (GR antagonist) injected I.P. at 10 mg/kg immediately after novel object recognition (NOR) training. **b.** % Time exploring the old and new objects (means ± SEM of n = 12 mice with NC *ad libitum*+vehicle, 11 with HF *ad libitum*+vehicle, 12 with NC *ad libitum*+RU486, 8 with HF *ad libitum*+RU486). Three-way ANOVA: Effect of RU486 *F*(1,39)=67 *p*<0.0001, effect of objects *F*(1,39) = 2.7 *p* = 0.1, effect of diet *F*(1,39) = 0 *p* = 0.9, effect of diet x objects *F*(1,39) = 0.9 *p* = 0.3, effect of objects X RU486 *F*(1,39) = 4.8 *p* = 0.03, effect of diet X RU486 *F*(1,39) = 0 *p* = 0.9, effect of diet X RU486 X objects *F*(1,39) = 0.7 *p* = 0.3. See **table S2** for post-hoc analysis \**p* = 0.01. HC: high fat/sugar diet, NC: normal chow. **c.** Remodelling of dendritic spines in *Thy1*-YFP males (n = 28) fed normal chow *ad libitum*. Arrows point to dynamic spines. Scale = 3 µm. **d.** % Spine gains and losses (means ± SEM) in S1 (n = 16 mice) and dCA1 (n = 12 mice). Two-way ANOVA: interaction of spine dynamics and RU486 *F*(1,20) = 14.9 *p* = 0.001 in S1. Kruskal-Wallis test *p* = 0.002 in CA1: Effect of RU486 on additions *p* = 0.003, eliminations *p* = 0.08. See **table S2** post-hoc analysis \*\**p* < 0.01. **e.** GR phosphorylation at sites dependent (S226) and independent (S134) of glucocorticoids in males (n = 30). **f.** Intensity (means ± SEM) of p-GR S134 in S1 (n = 11 mice) and dCA1 (n = 12 mice). Purple = TRF, black = *ad libitum*. Two-way ANOVA for S1 data: Effect of diet *F*(1,11) = 1 *p* = 0.3, TRF *F*(1,11) = 4.5 *p* = 0.05, interaction *F*(1,10) = 7.9 *p* = 0.01. For CA1 data: Effect of diet *F*(1,8) = 25 *p* = 0.001, TRF *F*(1,8) = 12.5 *p* = 0.007, interaction *F*(1,8) = 4.5 *p* = 0.06. See **table S2** for post-hoc analysis \**p* < 0.05. TRF: time restricted feeding, ad lib: ad libitum. **g.** Intensity (means ± SEM) of p-GR S226 in S1 (n = 12 mice) and dCA1 (n = 10 mice). Purple = TRF, black = *ad libitum*. Kruskal-Wallis test p = 0.7 for S1 data. Two-way ANOVA for CA1 data: Effect of diet *F*(1,10) = 0.8 *p* = 0.3, TRF *F*(1,10) = 0.6 *p* = 0.4, interaction *F*(1,10) = 0.01 *p* = 0.9. See **table S2** post-hoc analysis \**p* < 0.05.

### GR phosphorylation promotes memory retention by TRF on obesogenic food

We next used mouse carriers of S134A constitutive knock-in mutation (previously described (15)) to first determine the role of pS134-mediated signalling on the feeding-fasting cycle. Mutant mice had normal locomotor activity and anxiety despite an effect on body weight gain (interaction of genotype with diet and sex *p*=0.03 but not with TRF *p*=0.7, **Supplementary Fig. 6 and Table S2)**. Brains collected 1 hour after NOR recall showed lesser Fos-induction in S1 (**Fig. 6a**) and more in dCA1 (**Fig. 6b**) compared to WT controls, regardless of diet and meal scheduling, suggesting a lack of pS134 occluded the aversive effects of HFD and prevented the beneficial effects of TRF. Behaviourally, mutant mice exhibited NOR memory deficits early (**Fig. 6c**) that persisted on weeks 8 and 12 thereby mimicking the effect of HFD (**Fig. 6d**,**e**). Yet, TRF failed to correct memory deficits in HFD-fed mutant mice contrary to wildtype controls, suggesting that pS134 is required for recognition memory (**Fig. 6e**).

**Fig. 6.**
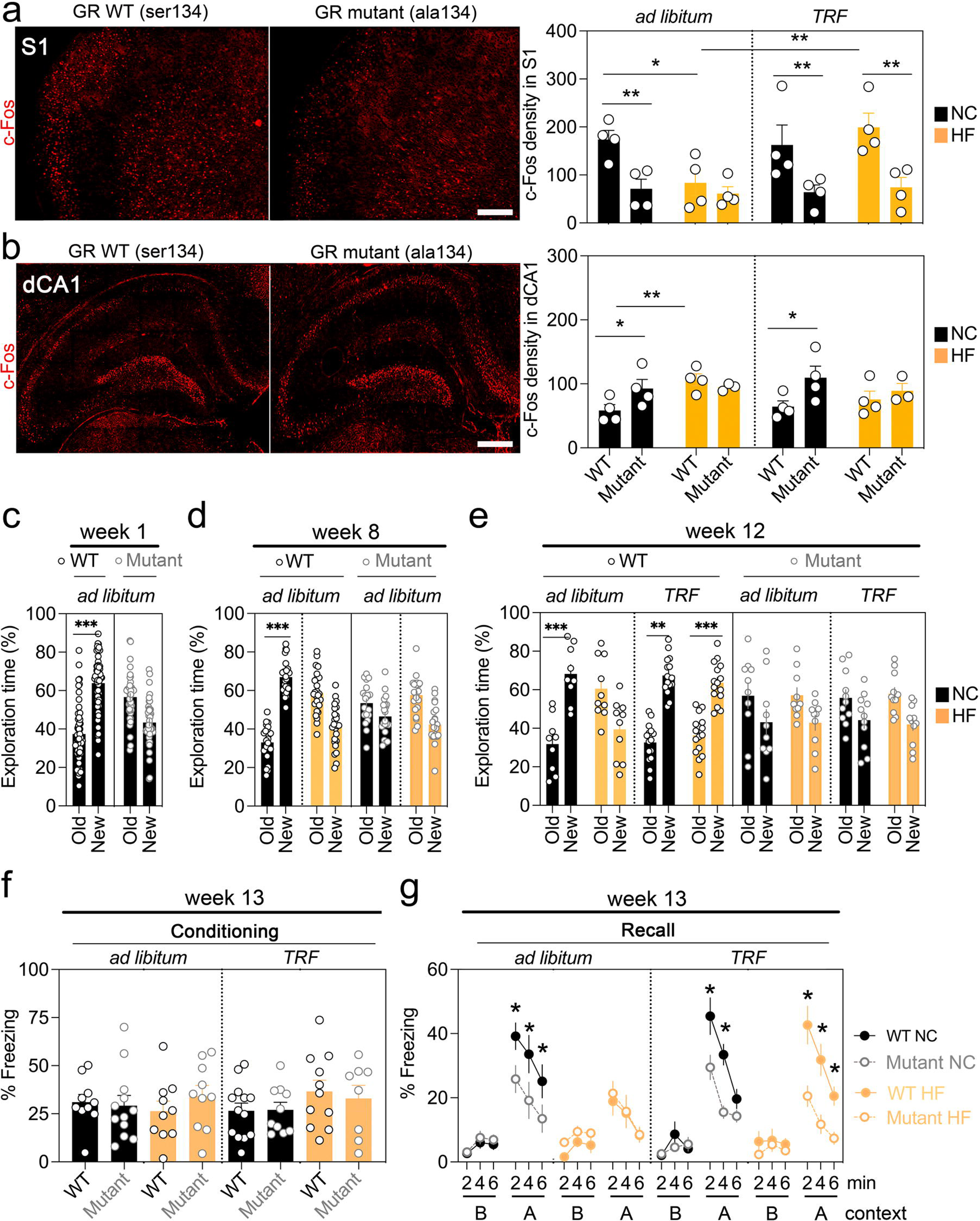
GR phosphorylation promotes memory retention by TRF in mice fed obesogenic food. **a.** Fos induction in S1 1 hour after object recognition memory test (means ± SEM, n = 16 WT GR^S134/S134^ and 16 mutants GR^A134/A134^ homozygotes). Three-way ANOVA: Effect of genotype *F*(1,24) = 24 *p* < 0.0001; interaction of diet and regimen *F*(1,24) = 4.3 *p* = 0.04 post-hoc analysis \**p* < 0.05, \*\**p* < 0.01. See **table S2** for details. HC: high fat/sugar diet, NC: normal chow, TRF: time restricted feeding. **b.** Fos induction in dCA1 1 hour after object recognition memory test (means ± SEM, n = 16 WT, 14 mutants). Three-way ANOVA: Effect of genotype *F*(1,22)=5.4 *p*=0.02; interaction of diet and genotype *F*(1,22)=5 *p*=0.03 post-hoc analysis \**p* < 0.05, \*\**p* < 0.01. See **table S2** for details. **c.** % Time exploring the old and new objects 24 hours after the training. Data are means ± SEM of n = 55 WT GR^S134/S134^ homozygotes (n = 24 males, 31 females) and 42 mutants GR^A134/A134^ homozygotes (n = 24 males, 18 females). Three-way ANOVA: Effect of sex *F*(1,54) = 0.1 *p* = 0.7, genotype *F*(1,54) = 0.1 *p* = 0.7, effect of objects *F*(1,54) = 17 *p* < 0.0001. Consolidated 2-way ANOVA without sex factor: Effect of genotype *F*(1,95) = 0.9 *p* = 0.3, effect of objects *F*(1,95) = 9.2 *p* = 0.003, interaction *F*(1,95) = 32 *p* < 0.0001. See **table S2** for post-hoc analyses \*\*\**p* < 0.001 and effect size. **d.** % Time exploring the old and new objects 24 hours after the training. Data are means ± SEM of n = 24 WT NC (n = 9 males, 15 females), 25 WT HF (n = 12 males, 13 females), 21 mutants NC (n = 13 males, 8 females), 21 mutants HF (n = 11 males, 10 females). Four-way ANOVA: Effect of sex *F*(1,166) = 0 *p* > 0.9, genotype *F*(1,166) = 0 *p* > 0.9, effect of objects *F*(1,166) = 0 *p* > 0.9, effect of diet *F*(1,166)= 0 *p* > 0.9, sex X objects *F*(1,166) = 2 *p* = 0.08, sex X genotype *F*(1,166) = 0 *p* > 0.9, sex X diet *F*(1,166) = 0 *p* > 0.9, genotype X objects *F*(1,166) = 38 *p* < 0.0001, objects X diet *F*(1,166) = 110 *p* < 0.0001, genotype X diet *F*(1,166) = 0 *p* > 0.9, sex X genotype X objects *F*(1,166) = 2 *p* = 0.1, sex X genotype X diet *F*(1,166) = 0 *p* > 0.9, sex X diet X objects *F*(1,166) = 0.08 *p* = 0.7, genotype X diet X objects *F*(1,166) = 52 *p* < 0.0001, sex X genotype X diet X objects *F*(1,166) = 0.1 *p* = 0.7. Consolidated 3-way ANOVA without sex factor: Effect of genotype *F*(1,87) = 86 *p* < 0.0001, effect of objects *F*(1,87) = 0.4 *p* = 0.5, effect of diet *F*(1,87)=1385 *p*<0.0001, genotype X objects *F*(1,87) = 21 *p* < 0.0001, objects X diet *F*(1,87) = 51 *p* < 0.0001, genotype X diet *F*(1,87) = 265 *p* < 0.0001, genotype X diet X objects *F*(1,87) = 27 *p* = 0.0003. See **table S2** for post-hoc analyses \*\*\**p* < 0.001 and effect size. **e.** % Time exploring the old and new objects 24 hours after the training. Data are means ± SEM of n = 9 WT NC *ad lib* (n = 3 males, 6 females), 15 WT NC TRF (n = 6 males, 9 females), 10 WT HF *ad lib* (n = 4 males, 6 females), 15 WT HF TRF (n = 7 males, 8 females), 10 mutants NC *ad lib* (n = 6 males, 4 females), 11 mutants NC TRF (n = 7 males, 4 females), 10 mutants HF *ad lib* (n = 4 males, 6 females), 11 mutants HF TRF (n = 7 males, 4 females). Five-way ANOVA: No main effects of single factors. Interaction of diet X objects *F*(1,108) = 80 *p* < 0.0001, genotype X object *F*(1,108) = 19 *p* < 0.0001, diet X schedule x object *F*(1,108) = 10 *p* = 0.001, diet X genotype X object *F*(1,108) = 30 *p* < 0.0001, diet X sex X schedule X object *F*(1,108) = 10 *p* = 0.01, diet X genotype X object X sex X schedule *F*(1,108) = 7.3 *p* = 0.008. See **table S2** for post-hoc analysis \*\**p* < 0.01, \*\*\**p* < 0.001 and effect size. **f.** No effect of genotype, TRF or diet on freezing behaviour during fear conditioning. Data are means ± SEM of n = 10 WT NC *ad lib* (n = 4 males, 6 females), 11 WT NC TRF (n = 6 males, 5 females), 10 WT HF *ad lib* (n = 4 males, 6 females), 13 WT HF TRF (n = 6 males, 7 females), 12 mutants NC *ad lib* (n = 6 males, 6 females), 10 mutants NC TRF (n = 6 males, 4 females), 10 mutants HF *ad lib* (n = 4 males, 6 females), 8 mutants HF TRF (n = 5 males, 3 females). ANOVA: No effect of main factors or interactions. See **table S2** for details. **g.** Effect of TRF % freezing behaviour 24 hours post-conditioning in context A (without shocks) and context B (with the unpaired tone). Data are means ± SEM of n = 10 WT NC *ad lib* (n = 4 males, 6 females), 11 WT NC TRF (n = 6 males, 5 females), 10 WT HF *ad lib* (n = 4 males, 6 females), 13 WT HF TRF (n = 6 males, 7 females), 12 mutants NC *ad lib* (n = 6 males, 6 females), 10 mutants NC TRF (n = 6 males, 4 females), 10 mutants HF *ad lib* (n = 4 males, 6 females), 8 mutants HF TRF (n = 5 males, 3 females). Friedman’s test *p* < 0.0001 indicates an effect of all factors with repeated measures (timepoints) validated with post-hoc Wilcoxon’s test. Kruskal-Wallis test shows no effect of sex with context recall *p* = 0.9 but with tone recall *p* = 0.008 although the effect size with Cohen’s *d* = 0.5 is neglectable. In contrast, there are significant effects of context with genotype *p* < 0.0001, with diet *p* = 0.0002 and with TRF *p* = 0.003. Pairwise group comparison indicates effects of genotypes, diet and TRF with

Finally, we tested emotional memory in mutant mice using contextual fear conditioning. Although acquisition was undistinguishable between genotypes (**Fig. 6f**), freezing behaviour was reduced in mutants specifically during recall of context (**Fig. 6g**). Again, TRF had no effect in the mutants, suggesting that pS134 is also required for emotional memory.

## Discussion

The present study shows that cortico-hippocampal activities necessary for remembering are uncoupled by obesogenic food consumed *ad libitum* but not on meal scheduling. Similar cortico-hippocampal mechanisms for memory retrieval previously involved the parietal cortex (PC) of mice and humans (32). Interestingly, not only do neuroimaging studies in PC-lesioned individuals revealed its active role in memory retrieval (32), but also in typically developing children where co-activation of PC-hippocampus axis is seen upon exposure to food (47). Chemogenetic manipulations in mice proved that cortical mechanisms are sufficient for remembering (48). We predict that cortical pathways regulate attentional states and stabilize behavioural representations in the hippocampus (49), which is disrupted by HFD.

### Strengths

Our longitudinal approach showed a reversal of behavioural and structural correlates of memory in the *same mice* after a change of feeding-fasting cycle, compared to feeding obesogenic food *ad libitum* around the clock. We report an excessive engagement of pyramidal neurons in dCA1 and disengagement in S1 with converging lines of evidence (Fos-induction, engram-trapping, dendritic spine maintenance). In agreement, Golgi-stained dCA1 neurons also showed higher densities of dendritic spines in adult mice fed HFD since adolescence (50) associated with enhanced hippocampal LTP (40). Additionally, TRF on HFD did not change overall dCA1 spine density either in Golgi-stained neurons (50) or in *Thy1*-YFP neurons. However, we noted a specific improvement in turnover of newly formed spines, in agreement with the normalization of LTP by TRF (24).

One striking observation is that HFD triggered bidirectional changes during peri-adolescence, persisting into adulthood. *Inhibiting* dCA1 hyperexcitation before retrieval corrected memory impairment in HFD-fed mice (39,41), consistent with previous findings on emotional memories (51). Thus, optimal dCA1 excitation is necessary to retrieve contextual memories. Conversely, chronic gain of excitation bilaterally throughout dCA1 previously caused severe amnesia (51), similar to HFD-induced aberrant activity and LTP in dCA1 (9,40,41). In S1, on the other hand, chemogenetic *activation* of pyramidal neurons only improved memories in HFD-fed mice, but not in controls. Together, it can be predicted that augmenting S1 activity could moderate dCA1 activity as previously shown with multi-unit recordings (52). HFD-fed mice showed uncoupling between a weakened S1 and hyperactivated dCA1. We confirmed that scaling activities between S1 and dCA1 is critical using cross-regional DREADD experiments, wherein partial inhibition of S1 and dCA1 was sufficient to remember despite HFD *ad libitum*. Conversely, partial excitation of both S1 and dCA1 failed to improve HFD-induced memory deficits. Thus, downscaling hyperactive dCA1 was both necessary and sufficient to improve long-term memory. This result is consistent with a previous study showing that the most stable populations of dCA1 pyramidal neurons -in terms of dendritic spine remodelling to sensory stimulation- are the ones engaged in neural representations of behaviours (53). Therefore, the issue with eating too rich a diet for too long might come down to how dCA1 activity is shaped by multiple convergent streams of information that participate in creating an integrated neuronal representation of exploratory behaviours (32).

Our findings in mice are also in agreement with co-activated hippocampus and sensory cortex in response to energy-dense food in neurotypical human subjects (54,55), a response that is uncoupled after a meal in obese children as compared to lean children (56). We showed that prolonged consumption of energy-dense food in mice triggered region-specific changes in neuroplasticity that biased glucocorticoid phospho-signalling. Increased levels of corticosterone was detectable after 12 weeks of HFD in rats, associated with enhanced synaptic depression in CA1 neurons (57). Pharmacological blockade with RU486 promoted net spine loss in S1 and gains in dCA1, perhaps due to the types of neurons and synapses involved (58). Inhibiting GR action with RU486 post-learning previously ameliorated object location memory in rats fed HFD for 1 week (59), contrary to what we find in mice fed HFD for 12 weeks, suggesting that GR signalling evolved between acute and chronic exposure to energy-dense food. For instance, HFD decreased BDNF levels in the cortex along with cognitive impairments (60,61), which could bias GR phospho-signalling toward the S226-dependent pathway. Here, we show for the first time that HFD suppressed GR signalling via the S134-dependent pathway while bolstering the S226-dependent pathway. Experiments in mice lacking S134 occluded the effects of feeding-fasting cycle, suggesting it is required for bi-directional experience- dependent synaptic plasticity (15,62). This is in agreement with HFD causing a circadian phase-shift in the expression of BDNF (63), GR (64) and corticosterone levels (65). Therefore, we propose that chronic HFD impeded the pro-neurotrophic signalling of glucocorticoids (19,58,66) in the cortico-hippocampal axis coupling attention with remembering. Promoting GR phosphorylation with inhibitors (or genetic invalidation) of the GR-phosphatase PP5 previously ameliorated adipogenesis and glucose tolerance induced by HFD (67,68) but the effects on brain co-morbidities are yet to be explored.

In a recent transcriptomic study, we showed that the beneficial effects of 14 hrs TRF in the peri-adolescence HFD model also involved thyroid hormones and astrocytic-mediated regulation of glutamatergic transmission (35). Cross-talk between thyroid and glucocorticoid signalling is well known, it remains to be investigated in this model to reinstate the cortico-hippocampal coupling that sustains long-term memory, and to confront obesity-related memory deficits.

## Limitations

The present set of observations do not exclude several alternate possibilities. For instance, it is possible that TRF could benefit all types of Western-style diets. It can also not be excluded that TRF benefits on cognition could involve GR in adipocytes and hepatocytes (69). Moreover, while we used the peri-adolescent HFD model because it is well characterized in terms of metabolic, neuroinflammatory and cognitive outcomes (25,59,70–72), TRF would also benefit other models such as when HFD started at adulthood or during gestation. For instance, gestational HFD caused protracted GR desensitization to a corticosterone challenge in adulthood that amplified neuroinflammation (73). Likewise, drug-induced GR desensitization during gestation worsened glucose tolerance and inflammation by HFD after birth (74). Thus, it cannot be excluded that TRF could benefit cognitive functioning in all age categories, nor predict lasting effects on shorter protocols nor after the cessation of TRF. So far, only one study assessed the effects of one-week TRF, which was sufficient to improve short-term working memory in HFD-fed adult mice, but the effect on long-term memory was not tested (24). The involvement of glucocorticoid-binding to the mineralocorticoid receptor has not been tested and therefore cannot be excluded from the regulation of dendritic spine remodelling, learning and memory (45,58,66). It cannot be excluded that cortico-hippocampal coupling is also necessary to retrieve remote memories (weeks/months after learning) by TRF. Despite the remodelling of dendritic spines by TRF in the cortico-hippocampal axis, it cannot be excluded that sprouting dendritic spines harbour silent synapses like in the stressed amygdala (75), which *in vivo* AMPA-uncaging could rule out as previously (38). Finally, the lack of reciprocal direct glutamatergic projections between S1 and dCA1 indicates that cortico-hippocampal coupling could result from polysynaptic circuits modulating dCA1 responses to sensory stimulation possibly via the dentate gyrus-CA3-CA1 pathway, the subiculum-dCA1 pathway and also through thalamic inputs (43). Future studies will advance the use of TRF beyond caveats and limitations.

## Contributors

Conceptualization: FJ, MPM, GF, EC; Methodology: YD, MAL, FJ, PC, AB, TSS; Investigations: YD, PC, EMA; Funding acquisition: MPM, GF; Project supervision: FJ; Writing: FJ; Editing: PC, FJ, EC, GF, MPM. All authors approved the final manuscript.

## Data sharing statement

DOI:10.5281/zenodo.10953723 without restriction.

## Declaration of interests

No conflict of interest.

## Notes

### Competing Interest Statement

The authors have declared no competing interest.

