## supplementary figures for "Feeding-fasting cycle of obesogenic food determines glucocorticoid neuromodulation of cortico-hippocampal activities sustaining long-term memory"

### Supplementary Methods

#### Study design

The protocol was not pre-registered.
Animals were randomly allocated to NC or HF ad lib groups.
No effect of sex was noted. So we pooled males and females, further distributed randomly to the following groups:

- NC TRF
- NC ad lib
- HF ad lib
- HF TRF

The outcome measures are as follows:

- object recognition memory
- freezing behaviour
- locomotor activity
- anxiety in the open field
- body weight
- dendritic spine addition
- elimination
- survival
- Cfos+ cell number
- Cfos+YFP+ cell number
- Cfos+Tomato+ cell number
- Tomato cell number
- pS134 levels
- pS226 levels
- GR levels.

Rationale for repeated measures as follows:

- Longitudinal studies to track impairment with HF and reversal with TRF.
- Chemogenetic to determine functional interaction of transgene and CNO. Multisite combinations to assess dominant effect between S1 and CA1.

The numbers of repeated measures is as follow:

- NOR behaviour was performed 3 times at week 1, 8, 12 post-weaning.
- Fear conditioning was performed once at week 13 post-weaning.
- Neuroimaging was performed 4 times in cortex at week 1, 8, 12, 13 post-weaning.
- Neuroimaging was performed 3 times in CA1 at week 8, 12, 13 post-weaning.
- RU486/vehicle injection was performed once at week 12 post-weaning.
- CNO/vehicle injection was performed once at week 12 post-weaning.

There are 13 experiments with the following group repartition:

- Experiment 1 has 4 groups: NC ad lib, NC TRF, HF ad lib, HF TRF to test object memory 3 times and emotional memory once.
- Experiment 2 has 4 groups: NC ad lib, NC TRF, HF ad lib, HF TRF for neuroimaging 4 times.

- Experiment 3 has 4 groups: NC ad lib, NC TRF, HF ad lib, HF TRF for post-mortem histology in thy1YFP.
- Experiment 4 has 4 groups: NC ad lib, NC TRF, HF ad lib, HF TRF for post-mortem histology in FosTRAP2;Ai14 mice for engram capture.
- Experiment 5 has 4 groups: NC ad lib +vehicle, NC ad lib +CNO, HF ad lib + vehicle, HF ad lib +CNO for DREADD-Gq in S1 on object memory.
- Experiment 6 has 4 groups: NC ad lib +vehicle, NC ad lib +CNO, HF ad lib + vehicle, HF ad lib +CNO for DREADD-Gi in CA1 on object memory.
- Experiment 7 has 6 groups: HF ad lib sham +vehicle, HF ad lib sham +CNO, HF ad lib Gi+vehicle, HF ad lib Gi+CNO, HF ad lib Gq+ vehicle, HF ad lib Gq+CNO for object memory.
- Experiment 8 has 4 groups: NC ad lib+vehicle, NC ad lib+RU486, HF ad lib+vehicle, HF ad lib+RU486 to test object memory.
- Experiment 9 has 4 groups: NC ad lib+vehicle, NC ad lib+RU486 imaged in S1 and NC ad lib+vehicle, NC ad lib+RU486 imaged in CA1.
- Experiment 10 has 4 groups: NC ad lib, NC TRF, HF ad lib, HF TRF for histology in S1 and CA1.
- Experiment 11 has 8 groups: WT NC ad lib, WT NC TRF, WT HF ad lib, WT HF TRF and mutant NC ad lib, mutant NC TRF, mutant HF ad lib, mutant HF TRF for histology in S1 and CA1.
- Experiment 12 has 8 groups: WT NC ad lib, WT NC TRF, WT HF ad lib, WT HF TRF and mutant NC ad lib, mutant NC TRF, mutant HF ad lib, mutant HF TRF for object memory.
- Experiment 13 has 8 groups: WT NC ad lib, WT NC TRF, WT HF ad lib, WT HF TRF and mutant NC ad lib, mutant NC TRF, mutant HF ad lib, mutant HF TRF for emotional memory.

**Animals**

All lines were maintained for more than 10 generations in C57BL6J background. All lines are backcrossed >10 generation in this background. All animals were bred and raised in the IGF experimental husbandry EOPS zone. Genotypes were determined before group allocation at P21. We used males and females from P21 to 4.5 months of age. GR phosphorylation numbering scheme by species is described elsewhere (57). All chemogenetic experiments are done in a mix of males and females as no sex difference were identified in the primary analysis (**Fig. 1 and supplementary Fig. S1**), and *in vivo* imaging was only done in males (due to excessive fluorescence in *Thy1-YFP* female mice). Mice are observed everyday by zootechnicians. Grid of endpoint measures includes body weight, behavior, posture, dehydration. All animals were handled by experimenters for a week before testing. All animals were habituated to the experiment-room for 1 hour before behavior tests. For surgeries, all animals were administered pre-op with lidocaine, anaesthetics and myorelaxant and post-op with meloxicam, carprofen.

**Open field**

Mice (males and females) positioned in the center freely explored an arena (50×50 cm, dim light ~50 lux) for 10 min. Animals were continuously recorded on video for offline scoring of locomotor activity by an observer blind for experimental groups. Total distance traveled and time spent in the center (29×29 cm) were determined with EzTrack (available on Github (58)). Data are presented in **supplementary Fig. S1**.

#### Novel object recognition

Mice (males and females) positioned in the center freely explored a L-shaped arena (30×10 cm, dim light ~50 lux) for 10 min on day 1 for habituation, with identical objects on each side on day 2, and with one old/previously explored (Lego blocks) and one novel object (falcon tube) on each side on day 3. Animals were continuously recorded on video for offline scoring of object exploration by an observer blind to experimental groups. In **Fig. 4**, we reported % time of object exploration as New/(New+Old) and Old/(New+Old). In **Fig. 1**, there was too many groups to compare (2 sexes, 3 time points, 2 diets, 2 feeding schedules). Therefore, we opted for the reporting of an object preference index calculated as (New-Old)/(New+Old). For the chemogenetic experiments (**Fig. 4**), the ligand CNO (clozapine-N-Oxide) 2 mg/kg dissolved in 0.5% DMSO in saline (used a vehicle control) was injected intra-peritoneally 45 min before the retrieval test on day 3. For the pharmacology experiments (**Fig. 5**), RU486 10 mg/kg dissolved in 2% ethanol in saline (used a vehicle control) was injected IP after the same objects exploration session to disrupt memory consolidation.

#### Contextual fear conditioning

This task was always done at the end because it is aversive and could influence the other behavioral metrics if done earlier. A week before testing, mice (males and females) were habituated to the experimenter. One day before conditioning, mice were left to explore without any stimulation for 3 min the neutral context B: rectangular opaque box without grid on the floor (18×35×22 cm) cleaned with 1% acetic acid lights OFF. This pre-exposure allowed the mice to acclimate and become familiar with the chamber (*neutral context*) used for the tone re-exposure test. On the next day, mice were placed in context A: transparent Plexiglas chamber (20×20×30 cm) with an electric grid floor, cleaned with 70% ethanol, lights ON and conditioned with 2 electrical foot-shocks (unconditional stimulus: US 0.4 mA, 1s) and 2 tones (conditional stimulus CS 65 dB, 1000 Hz, 15s) given pseudo-randomly after and before the shocks. More precisely, during the unpairing procedure, 100s after being placed in the chamber animals received a shock, then, after a 20s interval, a tone; finally, after a 30s delay, the same tone and the same shock spaced by a 30s interval were presented. After 20s, animals were returned to their home cage. As the tone is never followed by shock delivery, animals identify the conditioning context (set of static background contextual cues that constitutes the environment in which the conditioning takes place), and not the tone, as the right predictor of the shock (predictive context condition). On day 2, mice were re-exposed to the tone (65dB 2min) in context B. More precisely, first 2 min (no tone), next 2 min (tone), and last 2 min (no tone). Conditioned response to the tone is expressed by the percentage of freezing during the tone presentation. 2h later mice were re-exposed to the conditioning context A for 6 min without any stimulation (no tone, no shock). Freezing to the context was calculated as the percentage of the total time spent freezing during the successive three blocks of 2 min periods of the test. Freezing, defined as a lack of all movement except respiration, was used as an index of conditioned fear response. Animals were continuously recorded on video for offline second-by-second scoring of freezing by an observer blind to experimental groups.

#### Intra-hippocampal optical window

*Thy1*-YFP mice anesthetized with ketamine (0.075 mg/g) and xylazine (0.01 mg/g) were kept warm at 37°C, with pre-surgical lidocaine (10 mg, drops) on skull, and ophthalmic ointment applied on the eyes. For the cortical window, the scalp is sutured and topped with antibiotic cream to avoid infection between imaging sessions. For the intra-hippocampal window, dura was removed with forceps and cortex removed. The glass plug was glued to the skull and fixed with the custom head plate with dental cement. Adhesive plastic was kept over the window was protected from dust until imaging. Postoperative care consisted of 1 mg/kg meloxicam administration for 3 days.

**2-photon image acquisition**

Images were acquired in the somatosensory cortex S1 with a FVMPE RS two-photon microscope (Olympus, Hamburg, Germany) equipped with a 20X, numerical aperture NA1.0 water-immersion objective (Apocromat, Carl Zeiss) and an InSight X3 femtosecond-pulsed infrared laser (Spectra-Physics, Evry, France) for optimal fluorescence excitation and emission separation. Laser power was adjusted with the depth from 15 mW superficially and kept below 30 mW. For hippocampal imaging, the cover glass was rinsed with water and filled with ASCF. Images were acquired in dCA1 with the same microscope equipped with a long-range working distance (4 mm) objective 25x NA1.05 water-immersion (XLPLN25XWMP2, Olympus). Excitation was 960 nm for YFP. Images in S1 were taken with a digital zoom of 7.2x at each image session using 0.75 µm step with a scanning dwell time of 2.55 µsec per pixel. Images in dCA1 were taken with a digital zoom of 7.2x at each image session using 1 µm step with a scanning dwell time of 2.55 µsec per pixel. Laser power was adjusted with the depth from 20 mW superficially up to 60 mW. Each scan stacks consists of images at 512x512 pixels resolution.

**Image post-processing and analysis**

The field of view (200×200×150 µm) in consecutive images was realigned with RegStack plugin and distances between nearest spines along dendrites measured with ImageJ. Dendritic spines were marked manually with the Cell counter plugin of ImageJ. Two or more additions (or eliminations) of spines ≤5 µm along a dendrite define a dynamic cluster of formation (or elimination) as previously described (19). All clear headed-protrusions emanating laterally from the dendritic shaft were counted. In cortex, approximately 200 dendritic spines from at least 10-20 apical dendritic segments were counted per conditions throughout the imaging sessions and averaged per animal. We counted twice that amount in hippocampus given the higher spine density at apical dendrites. The presence, loss and gain of spines were counted between sessions for each segment and plotted as a function of distance to the nearest spine. Distance measurement between spines was set at the base of the neck to the base of the next spine following the trace of the dendritic shaft. The proportion of clustered formation (or elimination) equals the number of spines in clusters divided by the total number of new spines added (or eliminated) between imaging sessions. We used a bootstrapping method to ensure that spine clusters are not random as previously (39). New spine survival was defined as the number of spines gained between image 1 and 2, and further maintained in image 3, normalized by the number of spines present in the first time point. Old spine survival corresponds to the maintenance of pre-existing spines across time points divided by the total number of spines at the first time point.

**Histology**

Brains were harvested following transcardial perfusion with phosphate buffered saline (PBS) followed with 4% paraformaldehyde, and post-fixed for 24h at 4°C. Free-floating coronal sections of 40 μm obtained with a vibratome, were rinsed in PBS then blocked in 5% normal goat (or donkey) serum, PBS, 0.1% triton X-100 for 2h at 25°C. Primary antibodies, RFP (1:3000, Rockland), Iba1 (1:1000, Abcam), GFAP (1:1000, Merck), c-Fos 9F6 (1:1000, Cell signaling), p-GR S134 (1:1000, homemade (62)) p-GR S226 (1:1000, Gift from M. Garabedian, NYU USA (63)), GR P20 (1:400, Santa Cruz biotechnologies) were incubated for 2 days at 4°C and secondary antibodies (1:2000, ThermoFisher Scientific) for 2h at 25°C. Sections were washed in PBS, 0.1% Triton and mounted in Fluoromount (Sigma Aldrich). Images were acquired with an epifluorescence microscope (Imager Z1, Carl Zeiss), and the overlap between c-Fos and YFP cells or c-Fos and tdTomato cells counted with FIJI by an experimentalist blind to the groups.

**Supplementary Figures**

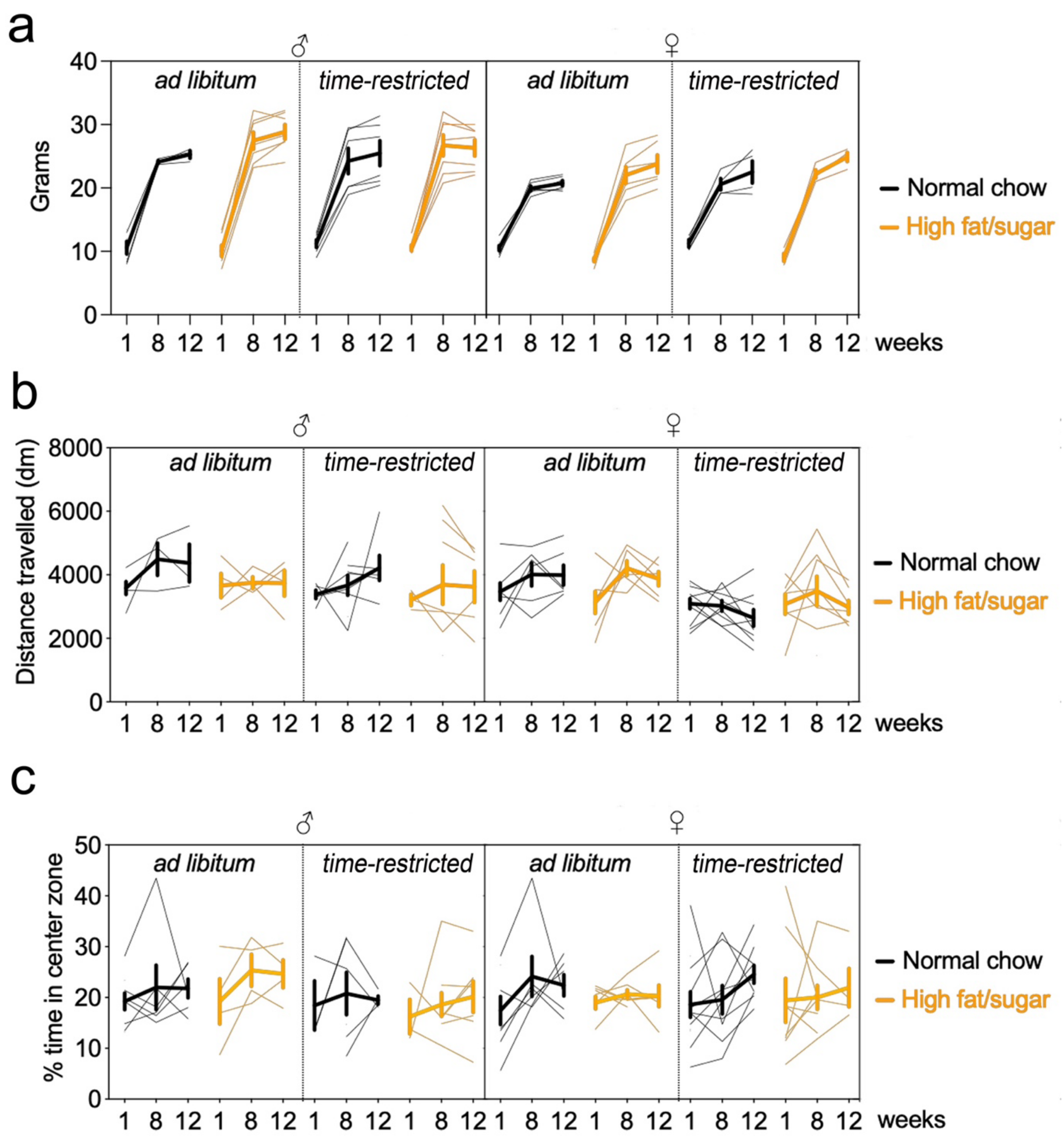


**Figure S1. No effect of diet nor schedule on locomotion, anxiety and body weight**
**a.** Developmental gain of body weight in males and females as a function of diet change (between week 1 and 8) and schedule change (between week 8 and 12). Bold lines indicate means ± SEM, thin lines individual subjects of n = 6 males + 6 females with NC ad libitum, 7 males + 7 females with HFS ad libitum, 6 males + 4 females with NC restricted, 7 males + 4 females with HFS restricted. Data has normal distribution (Shapiro-Wilk test $p = 0.52$). The use of the chi-square distribution instead of the F-distribution is justified by the lack of homoscedasticity (Levene's test $p < 0.0001$). Four-way ANOVA (model type III): Chisq analysis: Effect of sex $\chi2_{(1,35)} = 0.9$ $p = 0.3$, diet $\chi2_{(1,35)} = 1.2$ $p = 0.2$, TRF $\chi2_{(1,35)} = 0.07$ $p = 0.7$, **time**

**$\chi2_{(1,35)}$ = 292 *p* < 0.0001**, diet x sex $\chi2_{(1,35)}$ = 0.7 *p* = 0.3, sex x TRF $\chi2_{(1,35)}$ = 0.0001 *p* = 0.99, diet x TRF $\chi2_{(1,35)}$ = 0.02 *p* = 0.8, **sex x time $\chi2_{(2,70)}$ = 10 *p* = 0.005**, **diet x time $\chi2_{(2,70)}$ = 14 *p* = 0.0008**, TRF x time $\chi2_{(2,70)}$ = 0.3 *p* = 0.8, sex x diet x TRF $\chi2_{(1,35)}$ = 0.02 *p* = 0.8, sex x diet x time $\chi2_{(2,70)}$ = 0.5 *p* = 0.7, sex x time x TRF $\chi2_{(2,70)}$ = 3.5 *p* = 0.1, diet x TRF x time $\chi2_{(2,70)}$ = 0.01 *p* = 0.9, sex x diet x TRF x time $\chi2_{(2,70)}$ = 0.3 *p* = 0.8. Pairwise comparisons with post-hoc Tukey's test indicate no effect of sex within groups at each time points. There is no effect of single factors except time, which interacts with sex and diet.

**b.** Distance travelled in the open field in males and females as a function of diet change (between week 1 and 8) and schedule change (between week 8 and 12). Bold lines indicate means ± SEM, thin lines individual subjects of n = 6 males + 6 females with NC ad libitum, 4 males +7 females with HFS ad libitum, 6 males + 9 females with NC restricted, 6 males + 7 females with HFS restricted. Data has normal distribution (Shapiro-Wilk test *p* = 0.06) and homoscedasticity (Levene's test *p* = 0.13). Four-way ANOVA (model type III): Chisq analysis: Effect of sex $\chi2_{(1,56)}$ = 1.5 *p* = 0.2, diet $\chi2_{(1,56)}$ = 0.9 *p* = 0.3, TRF $\chi2_{(1,56)}$ = 0.006 *p* = 0.9, time $\chi2_{(1,56)}$ = 4.5 *p* = 0.1, diet x sex $\chi2_{(1,56)}$ = 0.5 *p* = 0.4, sex x TRF $\chi2_{(1,56)}$ = 0.6 *p* = 0.4, diet x TRF $\chi2_{(1,56)}$ = 0.5 *p* = 0.4, sex x time $\chi2_{(2,71)}$ = 1.4 *p* = 0.4, diet x time $\chi2_{(2,71)}$ = 0.6 *p* = 0.7, TRF x time $\chi2_{(2,71)}$ = 2.7 *p* = 0.2, sex x diet x TRF $\chi2_{(1,56)}$ = 0.7 *p* = 0.4, sex x diet x time $\chi2_{(2,71)}$ = 1.3 *p* = 0.5, sex x time x TRF $\chi2_{(2,71)}$ = 1.5 *p* = 0.4, diet x TRF x time $\chi2_{(2,71)}$ = 0.02 *p* = 0.9, sex x diet x TRF x time $\chi2_{(2,71)}$ = 0.6 *p* = 0.7. There is no effect of single factors nor interactions between factors.

**c.** % time spent in the center of the open field in males and females as a function of diet change (between week 1 and 8) and schedule change (between week 8 and 12). Bold lines indicate means ± SEM, thin lines individual subjects of n = 6 males + 6 females with NC ad libitum, 4 males + 7 females with HFS ad libitum, 6 males + 9 females with NC restricted, 7 males + 9 females with HFS restricted. Data has homogeneity of variance (Levene's test *p* = 0.1). The use of the chi-square distribution instead of the F-distribution is justified by the lack of normality (Shapiro-Wilk test *p* < 0.0001). Four-way ANOVA (model type III): Chisq analysis: Effect of sex $\chi2_{(1,42)}$ = 0.2 *p* = 0.6, diet $\chi2_{(1,42)}$ = 0.1 *p* = 0.7, TRF $\chi2_{(1,42)}$ = 0.01 *p* = 0.9, time $\chi2_{(2,72)}$ = 0.1 *p* = 0.9, diet x sex $\chi2_{(1,42)}$ = 0.4 *p* = 0.5, sex x TRF $\chi2_{(1,42)}$ = 0.01 *p* = 0.8, diet x TRF $\chi2_{(1,42)}$ = 0.01 *p* = 0.89, sex x time $\chi2_{(2,72)}$ = 1.6 *p* = 0.4, diet x time $\chi2_{(2,72)}$ = 1.1 *p* = 0.5, TRF x time $\chi2_{(2,72)}$ = 0.3 *p* = 0.8, sex x diet x TRF $\chi2_{(1,42)}$ = 0.03 *p* = 0.8, sex x diet x time $\chi2_{(2,72)}$ = 2.3 *p* = 0.3, sex x time x TRF $\chi2_{(2,72)}$ = 0.4 *p* = 0.7, diet x TRF x time $\chi2_{(2,72)}$ = 0.6 *p* = 0.7, sex x diet x TRF x time $\chi2_{(2,72)}$ = 0.1 *p* = 0.5. There is no effect of single factors nor interactions between factors.

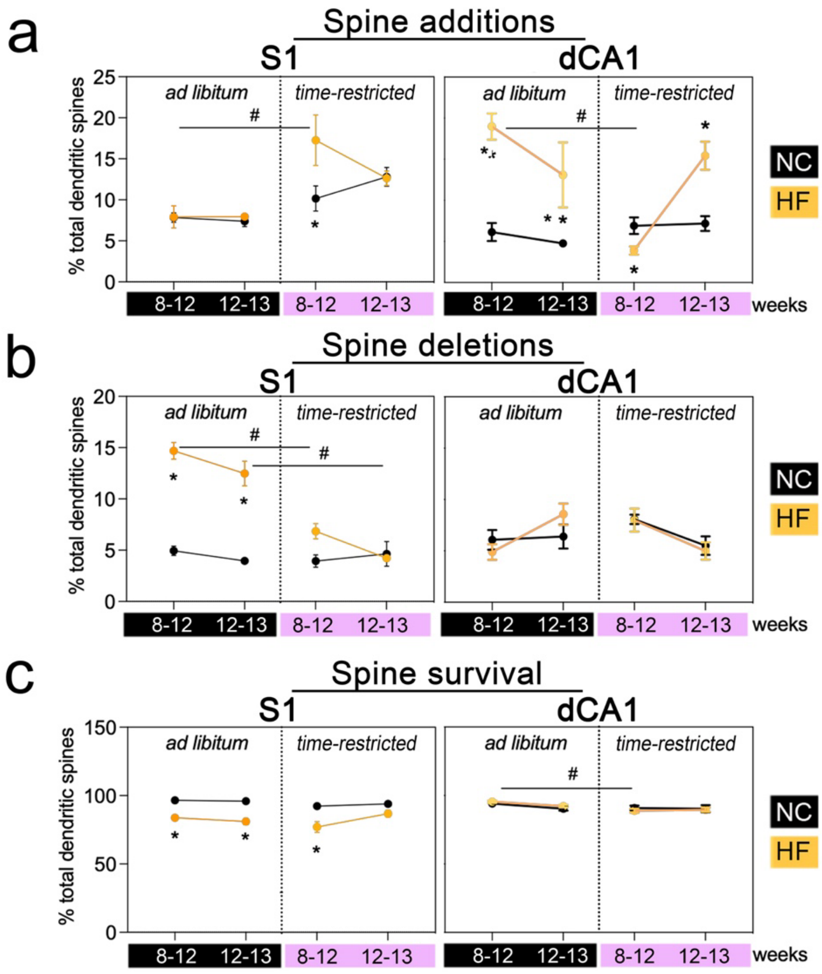


**Figure S2. TRF resets bi-directional spine remodeling in S1 and dCA1 of mice fed HFS food.**

**a.** % Spine additions in somatosensory cortex S1 and dorsal hippocampus subfield dCA1 between week 8 and 12, and between week 12 and 13 acquired in vivo by 2-photon microcopy in *Thy1*-YFP males only. Data are means ± SEM of n = 6 mice/group. Data has normal distribution (Shapiro-Wilk test $p > 0.05$) and homoscedasticity (Levene's test $p > 0.05$). Three-way ANOVA for *S1*: Effect of time $F_{(1,20)} = 0.2$ $p = 0.6$, schedule $F_{(1,20)} = 23$ $p < 0.0001$, diet $F_{(1,20)} = 3.1$ $p = 0.09$, time x schedule $F_{(1,20)} = 0.06$ $p = 0.8$, time x diet $F_{(1,20)} = 3.4$ $p = 0.08$, schedule x diet $F_{(1,20)} = 3.2$ $p = 0.08$, time x schedule x diet $F_{(1,20)} = 4.5$ $p = 0.04$. Post-hoc comparisons with Sidak test: NC restricted vs HFS restricted $^{*}p = 0.01$; HFS ad lib vs HFS restricted $^{\#}p = 0.005$. For *CA1*: Effect of time $F_{(1,20)} = 1.7$ $p = 0.2$, schedule $F_{(1,20)} = 5.4$ $p = 0.03$, diet $F_{(1,20)} = 46$ $p < 0.0001$, time x schedule $F_{(1,20)} = 28$ $p < 0.0001$, time x diet $F_{(1,20)} = 2$ $p = 0.1$, schedule x diet $F_{(1,20)} = 16$ $p = 0.0007$, time x schedule x diet $F_{(1,20)} = 17$ $p = 0.0004$. Post-hoc comparisons with Sidak test: NC ad lib vs HFS ad lib $^{*}p < 0.0001$ and $^{*}p = 0.0005$; NC restricted vs HFS restricted $^{*}p < 0.0001$; HFS ad lib vs HFS restricted $^{\#}p = 0.02$ and $^{\#}p < 0.0001$. HF: high fat/sugar diet, NC: normal chow. Black color: ad libitum, pink color: time restricted feeding.

**b.** % Spine eliminations in somatosensory cortex S1 and dorsal hippocampus subfield dCA1 between week 8 and 12, and between week 12 and 13 acquired in vivo by 2-photon microcopy in *Thy1*-YFP males only. Data are means ± SEM of n = 6 mice/group. Data has normal distribution (Shapiro-Wilk test $p > 0.05$) and homoscedasticity (Levene's test $p > 0.05$). Three-way ANOVA mixed-effect *for S1*: Effect of time $F_{(1,40)} = 8$ $p = 0.005$, schedule $F_{(1,40)} = 83$ $p < 0.0001$, diet $F_{(1,22)} = 143$ $p < 0.0001$, time x schedule $F_{(1,40)} = 1.9$ $p = 0.1$, time x diet $F_{(1,40)} = 5.3$ $p = 0.02$, schedule x diet $F_{(1,40)} = 91$ $p < 0.0001$, time x schedule x diet $F_{(1,40)} = 1.6$ $p = 0.2$. Post-hoc comparisons with Sidak test: NC ad lib vs HFS ad lib $^{*}p < 0.0001$; HFS ad lib vs HFS restricted $^{\#}p < 0.0001$. For *CA1*: Effect of time $F_{(1,20)} = 1.4$ $p = 0.2$, schedule $F_{(1,20)} = 0.5$ $p = 0.4$, diet $F_{(1,20)} = 0.3$ $p = 0.5$, time x schedule $F_{(1,20)} = 9.4$ $p = 0.006$, time x diet $F_{(1,20)} = 0.3$ $p = 0.5$, schedule x diet $F_{(1,20)} = 0.12$ $p = 0.7$, time x schedule x diet $F_{(1,20)} = 3.4$ $p = 0.07$. No difference in post-hoc comparisons with Sidak test. HF: high fat/sugar diet, NC: normal chow. Black color: ad libitum, pink color: time restricted feeding.

**c.** % Spine survival in somatosensory cortex S1 and dorsal hippocampus subfield dCA1 between week 8 and 12, and between week 12 and 13 acquired in vivo by 2-photon microcopy in *Thy1*-YFP males only. Data are means ± SEM of n = 6 mice/group. Data has normal distribution (Shapiro-Wilk test $p > 0.05$) and homoscedasticity (Levene's test $p > 0.05$). Three-way ANOVA *for S1*: Effect of time $F_{(1,40)} = 4.9$ $p = 0.03$, schedule $F_{(1,40)} = 1.7$ $p = 0.2$, diet $F_{(1,40)} = 73$ $p < 0.0001$, time x schedule $F_{(1,40)} = 6.9$ $p = 0.01$, time x diet $F_{(1,40)} = 2.5$ $p = 0.1$, schedule x diet $F_{(1,40)} = 1.5$ $p = 0.2$, time x schedule x diet $F_{(1,40)} = 4.1$ $p = 0.04$. Post-hoc comparisons with Sidak test: NC ad lib vs HFS ad lib $^{*}p < 0.001$; HFS ad lib vs HFS restricted $^{\#}p = 0.001$ and $^{\#}p = 0.04$. For *CA1*: Effect of time $F_{(1,20)} = 1.1$ $p = 0.2$, schedule $F_{(1,20)} = 9.8$ $p = 0.005$, diet $F_{(1,20)} = 0.02$ $p = 0.8$, time x schedule $F_{(1,20)} = 4.1$ $p = 0.055$, time x diet $F_{(1,20)} = 0.8$ $p = 0.3$, schedule x diet $F_{(1,20)} = 3.6$ $p = 0.07$, time x schedule x diet $F_{(1,20)} = 1.3$ $p = 0.2$. Post-hoc comparisons with Sidak test: HFS ad lib vs HFS restricted $^{\#}p = 0.002$. HF: high fat/sugar diet, NC: normal chow. Black color: ad libitum, pink color: time restricted feeding.

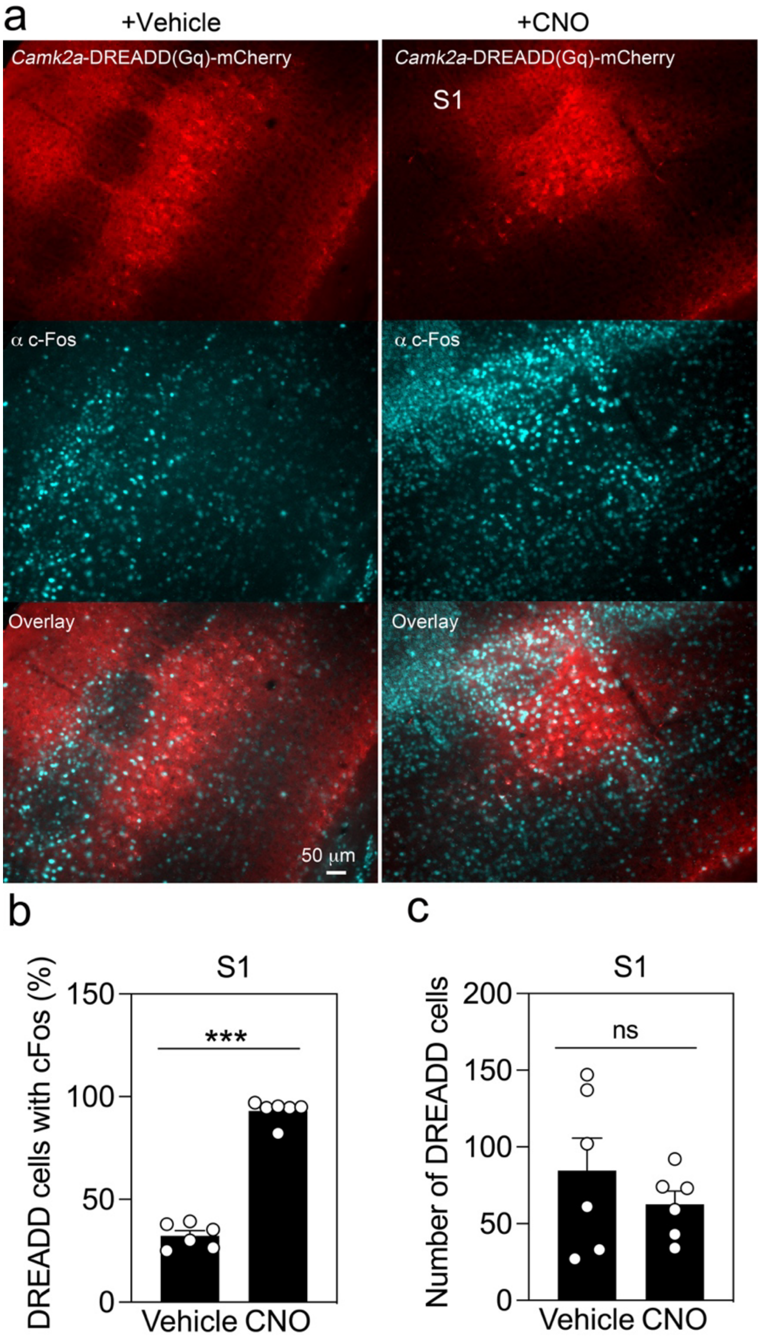


**Figure S3. Chemogenetic activation of pyramidal neurons in S1.**

**a.** Viral-mediated expression of *Camk2a*-DREADD(Gq)-mCherry in the somatosensory cortex S1 and cFos induction 45 min after I.P. injection of Clozapine-N-oxide (CNO as DREADD ligand) or vehicle as control.

**b.** % DREADD(Gq)-positive cells co-labelled with cFos antibodies. Data are means ± SEM (n = 6 males injected I.P. with vehicle, 6 males injected I.P. with CNO). Data has normal distribution (Shapiro-Wilk test $p > 0.05$) and homoscedasticity (Spearman's test $p > 0.05$). Statistical analysis done with 2-sided unpaired t-test $^{***}p < 0.0001$.

**c.** Number of DREADD(Gq)-positive cells. Data are means ± SEM (n = 6 males injected I.P. with vehicle, 6 males injected I.P. with CNO). Data has normal distribution (Shapiro-Wilk test

$p > 0.05$) and homoscedasticity (Spearman's test $p > 0.05$). Statistical analysis done with 2-sided unpaired t-test $p = 0.36$.

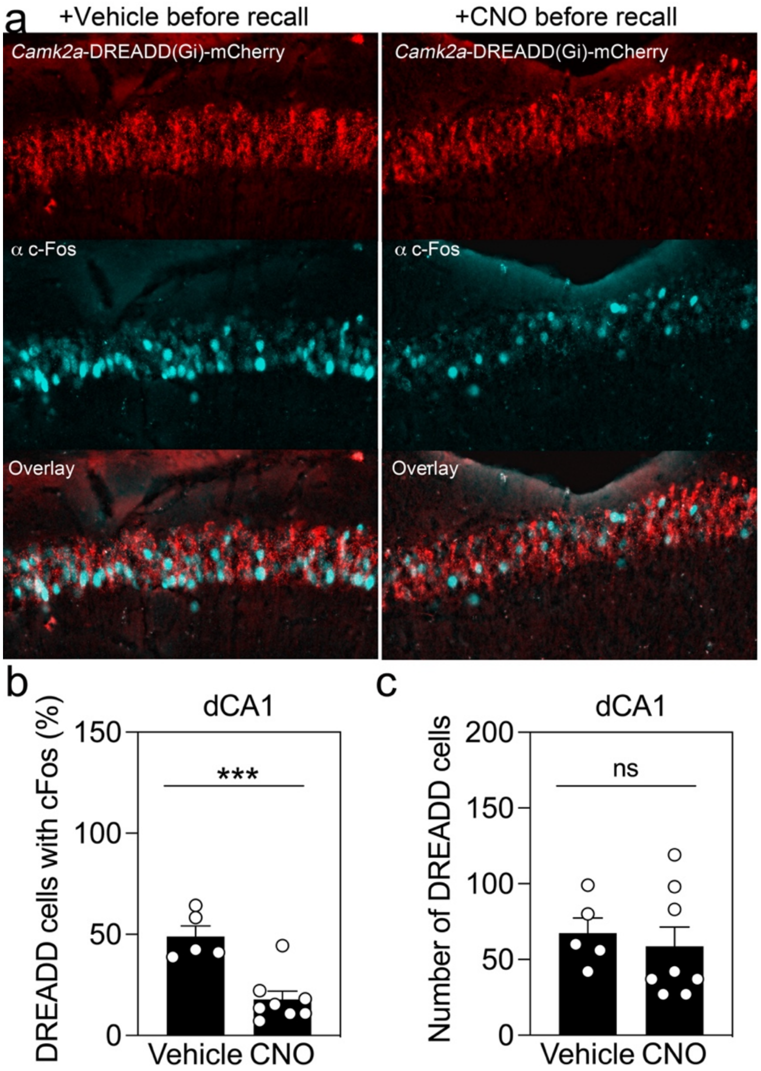


**Figure S4. Chemogenetic inhibition of pyramidal neurons in dCA1.**

**a.** Viral-mediated expression of *Camk2a*-DREADD(Gi)-mCherry in dorsal hippocampus subfield dCA1 and cFos induction 1 hr after object memory recall and I.P. injection of Clozapine-N-oxide (CNO) or vehicle as control.

**b.** % DREADD(Gq)-positive cells co-labelled with cFos antibodies. Data are means ± SEM (n = 5 males injected I.P. with vehicle, 8 males injected I.P. with CNO). Data has normal distribution (Shapiro-Wilk test $p > 0.05$) and homoscedasticity (Spearman's test $p > 0.05$). Statistical analysis done with 2-sided unpaired t-test $^{***}p = 0.0007$.

**c.** Number of DREADD(Gq)-positive cells. Data are means ± SEM (n = 5 males injected I.P. with vehicle, 8 males injected I.P. with CNO). Data has normal distribution (Shapiro-Wilk test $p > 0.05$) and homoscedasticity (Spearman's test $p > 0.05$). Statistical analysis done with 2-sided unpaired t-test $p = 0.64$.

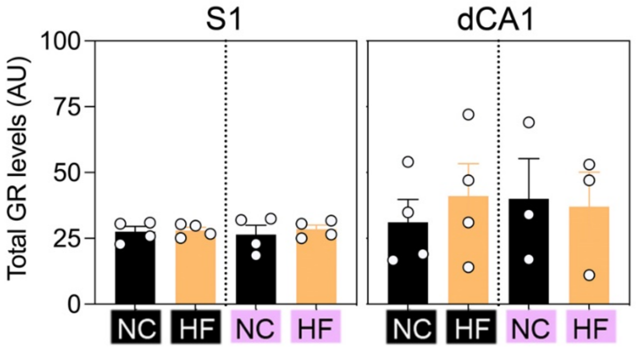


**Figure S5. No effect of diet and meal scheduling on GR levels in S1 and dCA1**
Intensity (arbitrary unit as AU) of GR immunoreactivity in S1 (n = 16) and dCA1 (n = 14) in male mice fed NC or HFS-diet either *ad libitum* or on scheduling (means ± SEM). Data has normal distribution (Shapiro-Wilk test $p > 0.05$) and homoscedasticity (Spearman's test $p > 0.05$). Three-way ANOVA: Effect of regions $F_{(1,22)} = 2.8$ $p = 0.1$, effect of regimen $F_{(1,22)} = 0.03$ $p = 0.8$, effect of diet $F_{(1,22)} = 0.16$ $p = 0.6$, effect of regimen x regions $F_{(1,22)} = 0.05$ $p = 0.8$, effect of regions x diet $F_{(1,22)} = 0.03$ $p = 0.8$, effect of regimen x diet $F_{(1,22)} = 0.2$ $p = 0.6$, effect of regimen x diet x regions $F_{(1,22)} = 0.3$ $p = 0.5$. HF: high fat/sugar diet, NC: normal chow. Black color: ad libitum, pink color: time restricted feeding.

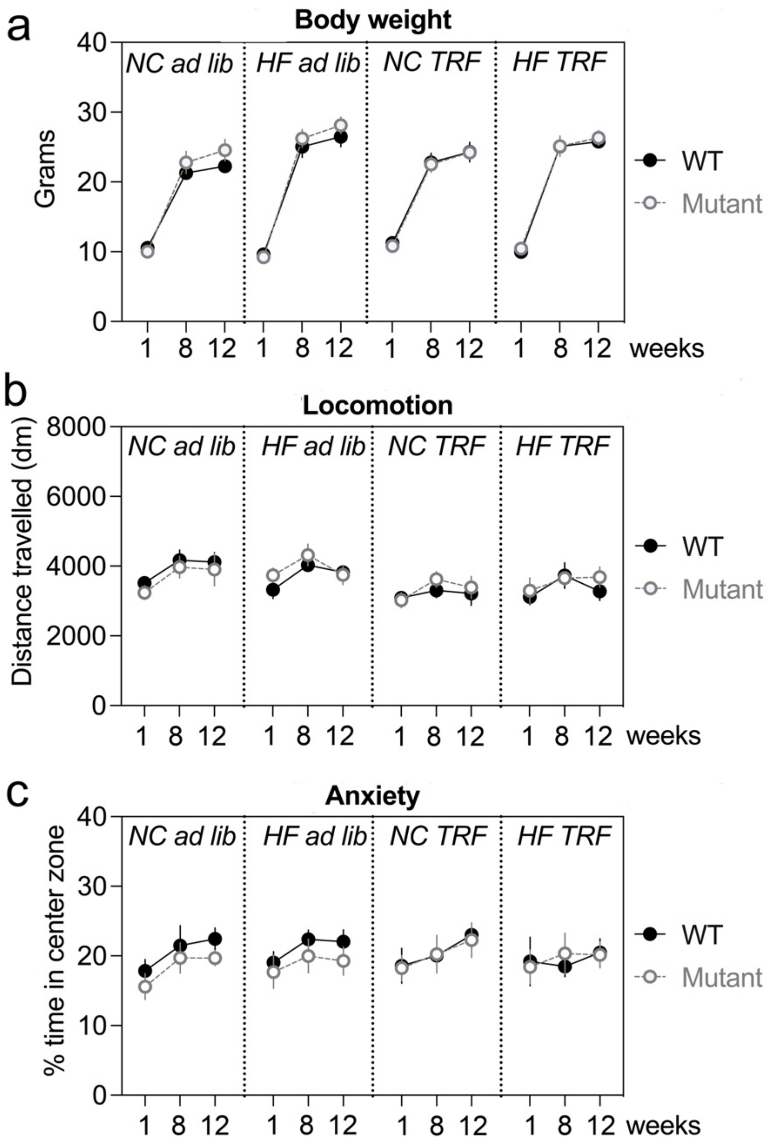


**Figure S6. No effect of genotype ($GR^{S134A}$) on locomotion, anxiety and body weight**

**a.** Developmental gain of body weight as a function of diet change (between week 1 and 8) and regimen change (between week 8 and 12). Data are means ± SEM. Repartition of 45 WT ($GR^{S134/S134}$) homozygotes is as follows: n = 6 males + 6 females with NC ad libitum, 7 males + 7 females with HF ad libitum, 6 males + 6 females with NC restricted, 7 males + 4 females with HF restricted. Repartition of 43 mutants ($GR^{A134/A134}$) homozygotes is as follows: n = 5 males + 5 females 5 KI males with NC ad libitum, 4 males + 6 females with HF ad libitum, 5 males + 4 females with NC restricted, 7 males + 6 females with HF restricted. Data has normal distribution (Shapiro-Wilk test $p = 0.47$). The use of the chi-square distribution instead of the F-distribution is justified by the lack of homoscedasticity (Levene's test $p < 0.0001$). Five-way ANOVA (model type III): Chisq analysis: Effect of sex $\chi2_{(1,68)} = 1.8$ $p = 0.1$, diet $\chi2_{(1,68)} = 0.9$ $p = 0.3$, TRF $\chi2_{(1,68)} = 0.3$ $p = 0.5$, **time $\chi2_{(1,68)} = 241$ $p < 0.0001$**, genotype $\chi2_{(1,68)} = 1$ $p = 0.3$, diet x sex $\chi2_{(1,68)} = 0.2$ $p = 0.5$, diet x TRF $\chi2_{(1,68)} = 0.1$ $p = 0.68$, **sex x time $\chi2_{(2,135)} = 36$ $p < 0.0001$**, diet x time $\chi2_{(2,135)} = 2.8$ $p = 0.2$, **TRF x time $\chi2_{(2,135)} = 7.5$ $p = 0.002$**, sex x genotype $\chi2_{(1,68)} = 2.8$ $p = 0.09$, diet x genotype $\chi2_{(1,68)} = 2.1$ $p = 0.1$, TRF x genotype $\chi2_{(1,68)} = 0.02$ $p = 0.1$, genotype x time $\chi2_{(2,135)} = 0.03$ $p = 0.9$, sex x diet x TRF $\chi2_{(1,68)} = 0.9$ $p = 0.3$, **sex x diet x time $\chi2_{(2,135)} = 9$ $p = 0.01$**, sex x TRF x time $\chi2_{(2,135)} = 1.6$ $p = 0.4$, diet x TRF x time $\chi2_{(2,135)} = 2.2$ $p = 0.3$, **sex x diet x genotype $\chi2_{(1,68)} = 4.5$ $p = 0.03$**, sex x TRF x genotype

$\chi2_{(1,68)}$ = 0.1 $p$ = 0.7, diet x genotype x TRF $\chi2_{(1,68)}$ = 0.03 $p$ = 0.8, **sex x genotype x time $\chi2_{(2,135)}$ = 7 $p$ = 0.02**, **sex x diet x time $\chi2_{(2,135)}$ = 9 $p$ = 0.01**, sex x TRF x time $\chi2_{(2,135)}$ = 1.6 $p$ = 0.4, diet x TRF x time $\chi2_{(2,135)}$ = 2.2 $p$ = 0.3, **sex x diet x genotype $\chi2_{(1,68)}$ = 4.5 $p$ = 0.03**, diet x time x genotype $\chi2_{(2,135)}$ = 1.3 $p$ = 0.5, time x genotype x TRF $\chi2_{(2,135)}$ = 3.2 $p$ = 0.1, sex x diet x TRF x time $\chi2_{(2,135)}$ = 0.09 $p$ = 0.9, sex x TRF x time x genotype $\chi2_{(2,135)}$ = 0.6 $p$ = 0.4, **sex x time x genotype x diet $\chi2_{(2,135)}$ = 6.7 $p$ = 0.03**, sex x TRF x time x genotype $\chi2_{(2,135)}$ = 0.1 $p$ = 0.9, TRF x diet x time x genotype $\chi2_{(2,135)}$ = 0.9 $p$ = 0.6, sex x TRF x diet x time x genotype $\chi2_{(2,135)}$ = 0.3 $p$ = 0.8. There is no effect of single factors except time. There is an interaction of sex with time, with time and genotype, diet and time, and with diet+time+genotype. The effect of genotype is only significant when interacting with the sex factor. Pairwise comparisons with post-hoc Tukey's test indicated no significant effect of sex between groups at each time points. HF: high fat/sugar diet, NC: normal chow. Ad lib: ad libitum, TRF: time restricted feeding.

**b.** Distance travelled in the open field in males and females as a function of diet change (between week 1 and 8) and regimen change (between week 8 and 12). Data are means ± SEM. Repartition of 45 WT ($GR^{S134/S134}$) homozygotes is as follows: n = 6 males + 6 females with NC ad libitum, 7 males + 7 females with HF ad libitum, 6 males + 6 females with NC restricted, 7 males + 4 females with HF restricted. Repartition of 43 mutants ($GR^{A134/A134}$) homozygotes is as follows: n = 5 males +5 females 5 KI males with NC ad libitum, 4 males + 6 females with HF ad libitum, 5 males + 4 females with NC restricted, 7 males + 6 females with HF restricted. Data has normal distribution (Shapiro-Wilk test $p$ = 0.07). The use of the chi-square distribution instead of the F-distribution is justified by the lack of homoscedasticity (Levene's test $p$ = 0.02). Five-way ANOVA (model type III): Chisq analysis: Effect of sex $\chi2_{(1,78)}$ = 0.01 $p$ = 0.8, diet $\chi2_{(1,93)}$ = 2.1 $p$ = 0.1, TRF $\chi2_{(1,93)}$ = 0.4 $p$ = 0.5, time $\chi2_{(1,93)}$ = 2.4 $p$ = 0.29, genotype $\chi2_{(1,93)}$ = 1 $p$ = 0.3, diet x sex $\chi2_{(1,93)}$ = 0.5 $p$ = 0.4, sex x TRF $\chi2_{(1,93)}$ = 0.01 $p$ = 0.8, diet x TRF $\chi2_{(1,93)}$ = 0.2 $p$ = 0.6, sex x time $\chi2_{(2,129)}$ = 0.2 $p$ = 0.8, diet x time $\chi2_{(2,129)}$ = 1.8 $p$ = 0.3, TRF x time $\chi2_{(2,129)}$ = 1.5 $p$ = 0.9, sex x genotype $\chi2_{(1,93)}$ = 0.28 $p$ = 0.59, diet x genotype $\chi2_{(1,93)}$ = 2.5 $p$ = 0.1, TRF x genotype $\chi2_{(1,93)}$ = 0.09 $p$ = 0.7, genotype x time $\chi2_{(2,129)}$ = 1.2 $p$ = 0.5, sex x diet x TRF $\chi2_{(1,93)}$ = 0.1 $p$ = 0.6, sex x diet x time $\chi2_{(2,129)}$ = 0.8 $p$ = 0.6, sex x TRF x time $\chi2_{(2,129)}$ = 0.4 $p$ = 0.8, diet x TRF x time $\chi2_{(2,129)}$ = 0.9 $p$ = 0.6, sex x diet x genotype $\chi2_{(1,93)}$ = 0.9 $p$ = 0.3, sex x TRF x genotype $\chi2_{(1,93)}$ = 0.001 $p$ = 0.9, diet x genotype x TRF $\chi2_{(1,93)}$ = 0.4 $p$ = 0.5, sex x genotype x time $\chi2_{(2,129)}$ = 1.2 $p$ = 0.5, genotype x diet x time $\chi2_{(2,129)}$ = 1.8 $p$ = 0.4, genotype x TRF x time $\chi2_{(2,129)}$ = 0.5 $p$ = 0.7, sex x diet x TRF x time $\chi2_{(2,129)}$ = 0.1 $p$ = 0.9, sex x diet x TRF x genotype $\chi2_{(1,93)}$ = 0.6 $p$ = 0.6, sex x time x genotype x diet $\chi2_{(2,129)}$ = 2.4 $p$ = 0.3, sex x TRF x time x genotype $\chi2_{(2,129)}$ = 0.2 $p$ = 0.9, TRF x diet x time x genotype $\chi2_{(2,129)}$ = 0.6 $p$ = 0.7, sex x TRF x diet x time x genotype $\chi2_{(2,129)}$ = 0.04 $p$ = 0.9. There is no effect of single factors nor interactions between factors. HF: high fat/sugar diet, NC: normal chow. Ad lib: ad libitum, TRF: time restricted feeding.

**c.** % time spent in the center of the open field in males and females as a function of diet change (between week 1 and 8) and regimen change (between week 8 and 12). Data are means ± SEM. Repartition of 45 WT ($GR^{S134/S134}$) homozygotes is as follows: n = 6 males + 6 females with NC ad libitum, 7 males + 7 females with HF ad libitum, 6 males + 6 females with NC restricted, 7 males + 4 females with HF restricted. Repartition of 43 mutants ($GR^{A134/A134}$) homozygotes is as follows: n = 5 males + 5 females 5 KI males with NC ad libitum, 4 males + 6 females with HF ad libitum, 5 males + 4 females with NC restricted, 7 males + 6 females with HF restricted. Data has homogeneity of variance (Levene's test $p$ = 0.24). The use of the chi-square distribution instead of the F-distribution is justified by the lack of normality (Shapiro-Wilk test

$p < 0.0001$). Five-way ANOVA (model type III): Chisq analysis: Effect of sex $\chi2_{(1,78)} = 0.7$ $p = 0.3$, diet $\chi2_{(1,78)} = 0.6$ $p = 0.4$, TRF $\chi2_{(1,78)} = 0.1$ $p = 0.7$, time $\chi2_{(1,78)} = 1.1$ $p = 0.5$, genotype $\chi2_{(1,78)} = 0.4$ $p = 0.4$, diet x sex $\chi2_{(1,78)} = 0.05$ $p = 0.8$, sex x TRF $\chi2_{(1,78)} = 0.2$ $p = 0.6$, diet x TRF $\chi2_{(1,78)} = 0.2$ $p = 0.6$, sex x time $\chi2_{(2,110)} = 2$ $p = 0.3$, diet x time $\chi2_{(2,110)} = 1$ $p = 0.5$, TRF x time $\chi2_{(2,110)} = 0.02$ $p = 0.9$, sex x genotype $\chi2_{(1,110)} = 0.9$ $p = 0.3$, diet x genotype $\chi2_{(1,78)} = 0.15$ $p = 0.6$, TRF x genotype $\chi2_{(1,78)} = 0.04$ $p = 0.8$, genotype x time $\chi2_{(2,110)} = 0.4$ $p = 0.8$, sex x diet x TRF $\chi2_{(1,78)} = 0.05$ $p = 0.8$, sex x diet x time $\chi2_{(2,110)} = 1$ $p = 0.6$, sex x TRF x time $\chi2_{(2,110)} = 0.4$ $p = 0.7$, diet x TRF x time $\chi2_{(2,110)} = 0.3$ $p = 0.8$, sex x diet x genotype $\chi2_{(1,78)} = 0.08$ $p = 0.7$, sex x TRF x genotype $\chi2_{(1,78)} = 0.07$ $p = 0.7$, diet x genotype x TRF $\chi2_{(1,78)} = 0.09$ $p = 0.7$, sex x genotype x time $\chi2_{(2,110)} = 2.5$ $p = 0.2$, genotype x diet x time $\chi2_{(2,110)} = 0.01$ $p = 0.9$, genotype x TRF x time $\chi2_{(2,110)} = 0.06$ $p = 0.9$, sex x diet x TRF x time $\chi2_{(2,110)} = 0.3$ $p = 0.8$, sex x diet x TRF x genotype $\chi2_{(1,78)} = 0.09$ $p = 0.7$, sex x time x genotype x diet $\chi2_{(2,110)} = 0.6$ $p = 0.7$, sex x TRF x time x genotype $\chi2_{(2,110)} = 0.8$ $p = 0.6$, TRF x diet x time x genotype $\chi2_{(2,110)} = 0.1$ $p = 0.9$, sex x TRF x diet x time x genotype $\chi2_{(2,110)} = 0.3$ $p = 0.8$. There is no effect of single factors nor interactions between factors. HF: high fat/sugar diet, NC: normal chow. Ad lib: ad libitum, TRF: time restricted feeding.
